# An increase in serial sarcomere number induced via weighted downhill running improves work loop performance in the rat soleus

**DOI:** 10.1101/2022.02.18.481073

**Authors:** Avery Hinks, Kaitlyn Jacob, Parastoo Mashouri, Kyle D. Medak, Martino V. Franchi, David C. Wright, Stephen H. M. Brown, Geoffrey A. Power

## Abstract

Increased serial sarcomere number (SSN) has been observed in rats via downhill running training due to the emphasis on active lengthening contractions; however, little is known about the influence on dynamic contractile function. Therefore, we employed 4 weeks of weighted downhill running training in rats, then assessed soleus SSN and work loop performance. We hypothesized trained rats would produce greater net work output during faster, higher-strain work loops due to a greater SSN. Thirty-one Sprague-Dawley rats were assigned to a control or training group. Weight was added during downhill running via a custom-made vest, progressing from 5-15% body mass. Following sacrifice, the soleus was dissected, and a force-length relationship was constructed. Work loops (active shortening followed by passive lengthening) were then performed about optimal muscle length (L_O_) at 1.5-3-Hz cycle frequencies and 1-7-mm strains to assess net work output. Muscles were then fixed in formalin at L_O_. Fascicle lengths and sarcomere lengths were measured and used to calculate SSN. Intramuscular collagen content and crosslinking were quantified via a hydroxyproline content and pepsin-solubility assay. Trained rats had longer fascicle lengths (+13%), greater SSN (+8%), greater specific active forces (+50%), and lower passive forces (–45-62%) than controls (P<0.05). There were no differences in collagen parameters (P>0.05). Net work output was greater (+101-424%) in trained than control rats for the 1.5-Hz loops at 1, 3, and 5-mm strains (*P*<0.05) and showed relationships with fascicle length (R^2^=0.14-0.24, *P*<0.05). These results suggest training-induced longitudinal muscle growth may improve dynamic performance.

## Introduction

Skeletal muscle remodels and adapts in response to specific conditions, as observed across the hierarchy of muscle structural organization (Gans and Bock, 1965; Gans and de Vree, 1987; Jorgenson et al., 2020). An example is longitudinal skeletal muscle growth, seen as muscle fascicle elongation and a corresponding increase in serial sarcomere number (SSN), termed sarcomerogenesis, following downhill running training due to the emphasis on active lengthening (eccentric) contractions (Butterfield et al., 2005a; Chen et al., 2020; Lynn et al., 1998). During unaccustomed eccentric exercise, sarcomeres are overstretched, resulting in suboptimal actin-myosin overlap. The most supported hypothesis for sarcomerogenesis following eccentric training is that an increase in SSN occurs to re-establish optimal crossbridge overlap regions (Butterfield et al., 2005a; Herring et al., 1984). While the influence of increased SSN on isometric contractile function is well-established (Lynn et al., 1998; Williams and Goldspink, 1978), less is known about the impact on measures of dynamic contractile function.

A physiologically relevant assessment of dynamic performance *in vitro* is the ‘work loop’: sinusoidal cycles of muscle shortening and lengthening with phasic bursts of stimulation meant to simulate locomotion (Josephson and Stokes, 1989; Schaeffer and Lindstedt, 2013). Work loops can incorporate a range of cycle frequencies (i.e., speeds of shortening/lengthening) and strains (i.e., muscle length changes), and are therefore influenced by force-velocity and force-length properties of muscle (Josephson, 1999). Hence, there are optimal strains and cycle frequencies for work loop performance, wherein active tension is as high as possible during shortening and as low as possible during subsequent lengthening (i.e., generates greater net work output) (Josephson and Stokes, 1989; Swoap et al., 1997). Sarcomerogenesis may result in placing sarcomeres closer to optimal length throughout the range of motion (especially at longer muscle lengths), thus allowing individual sarcomeres to shorten slower, optimizing force production across a wider range of muscle lengths based on the force-length and force-velocity relationships (Akagi et al., 2020; Drazan et al., 2019). SSN is also proportional to maximum shortening velocity (Wickiewicz et al., 1984). Taken together, sarcomerogenesis may improve work loop performance particularly in cycles with longer strains and faster cycle frequencies.

Contrary to the above hypothesis, Cox et al. (2000) increased SSN of the rabbit latissimus dorsi via 3 weeks of incremental surgical stretch and observed a decrease in maximum work loop power output compared to controls. They attributed this impaired work loop performance despite increased SSN to increased intramuscular collagen content caused by the surgical stretch. Increased collagen content and crosslinking enhance muscle passive stiffness (Brashear et al., 2020; Huijing, 1999; Kjær, 2004; Turrina et al., 2013), which can increase the work of passive lengthening and thereby decrease net work output. While collagen content and mRNAs and growth factors related to collagen synthesis can increase in the rat medial gastrocnemius and quadriceps femoris following isometric, concentric, and eccentric-based training (Han et al., 1999; Heinemeier et al., 2007; Zimmerman et al., 1993), collagen content and key enzymes related to collagen synthesis do not change in the rat soleus following short-term downhill running (Han et al., 1999) or long-term running training (Zimmerman et al., 1993). Collagen crosslinking was also shown to not change in the rat soleus with long-term running training (Zimmerman et al., 1993). Hence, the rat soleus in downhill running training may be a good model to assess the influence of sarcomerogenesis on work loop performance in the absence of increased collagen content.

Previous studies have employed downhill running in rodents with primarily an endurance training stimulus: on consecutive days with no progressive weighted overload (Butterfield et al., 2005a; Chen et al., 2020). A stronger stimulus provided by progressively loaded training may have a more pronounced impact on muscle architecture and mechanical function (Butterfield and Herzog, 2006; Farup et al., 2012). Furthermore, running on many consecutive days may minimize time for recovery and hence remodelling between exercise bouts (Hyldahl and Hubal, 2014). This distinction in training stimuli was offered previously as explanation for observations of only small adaptations in rat soleus SSN and mechanical function following downhill running training (Chen et al., 2020). Larger magnitude changes in muscle architecture and strength in animals following 3 days/week training programs support this perspective as well (Butterfield and Herzog, 2006; Ochi et al., 2007; Zhu et al., 2021).

To enhance the effect of downhill running training on muscle architecture, the present study employed downhill running 3 days/week and progressively increased the eccentric stimulus during running via a novel model incorporating weighted vests. The purposes were to: 1) assess how soleus SSN and intramuscular collagen adapt to eccentric training; and 2) investigate the influence on work loop performance. We hypothesized that soleus SSN would increase, and collagen content would not change following training. We also hypothesized that, due to training-induced sarcomerogenesis, work loop performance would improve, particularly in work loops with longer strains and faster cycle frequencies.

## Methods

### Animals

Thirty-one male Sprague-Dawley rats (sacrificial age ∼18 weeks) were obtained (Charles River Laboratories, Senneville, QC, Canada) with approval from the University of Guelph’s Animal Care Committee and all protocols following CCAC guidelines. Rats were housed at 23°C in groups of three and given ad-libitum access to a Teklad global 18% protein rodent diet (Envigo, Huntington, Cambs., UK) and room-temperature water. After a week of acclimation to housing conditions and familiarization with the vests and treadmills, rats were assigned to 1 of 2 groups: control (n = 18) or training (n = 13). Training consisted of 4 weeks of weighted downhill running 3 days/week. Approximately 72 hours following recovery from the final training day, rats were sacrificed via CO_2_ asphyxiation and cervical dislocation (Chen et al., 2020). We then immediately dissected the soleus and proceeded with mechanical testing. The soleus was chosen for this study due to its simple fusiform structure with fascicles running tendon to tendon (Williams and Goldspink, 1973), its expected lack of changes in collagen content and crosslinking (Han et al., 1999; Zimmerman et al., 1993), and its suitability for prolonged work loop experiments, being a primarily slow-fibered muscle (Caiozzo and Baldwin, 1997; Swoap et al., 1997). Due to measurement errors in characterizing the force-length relationship after determining L_O_, 2 rats (1 control, 1 training) were excluded from analysis of the force-length relationship. Similarly, due to a faulty setup of work loop protocols, the same 2 rats plus 1 other (2 control, 1 training) were excluded from work loop data analysis.

### Weighted vests

After extensive piloting, we designed a custom-made weighted rodent vest that is both well tolerated by the rat and sufficiently adds weight during running (Figure 1). A child-sized sock was cut into the shape of a vest: the end of the sock was cut off such that the ankle end went around the rat’s neck and the toe end went around the rat’s torso, then arm holes were cut on each side just distal to the neck hole. A T-shaped piece of foam paper was then super-glued on the back of the vest starting above the arm holes. A piece of Velcro was then placed across the wider section of the T-shaped foam paper. The apparatus for holding the weights was then constructed. A strip of packing tape was cut to be the same length as the wider section of the T-shaped foam paper. To allow for easier manipulation of the weight apparatus, a pipe cleaner of the same length was then cut and stuck across the midline of the tape, then a thinner piece of tape was placed over top to hold the pipe cleaner in place. Two 2 cm × 2 cm Ziplock bags were then stuck on the strip of tape over top of the pipe cleaner, equidistant from the middle. Additional small strips of tape were added as needed to secure these bags in place. A connecting piece of Velcro was then placed on the side of the tape opposite the bags, allowing it to be fastened to the T-shaped foam paper. The appropriate weights (various sizes of coins) were placed in the Ziplock bags. The bags were small enough to hold the coins securely in place, such that they did not move around as the rat ran.

**Figure 1:**
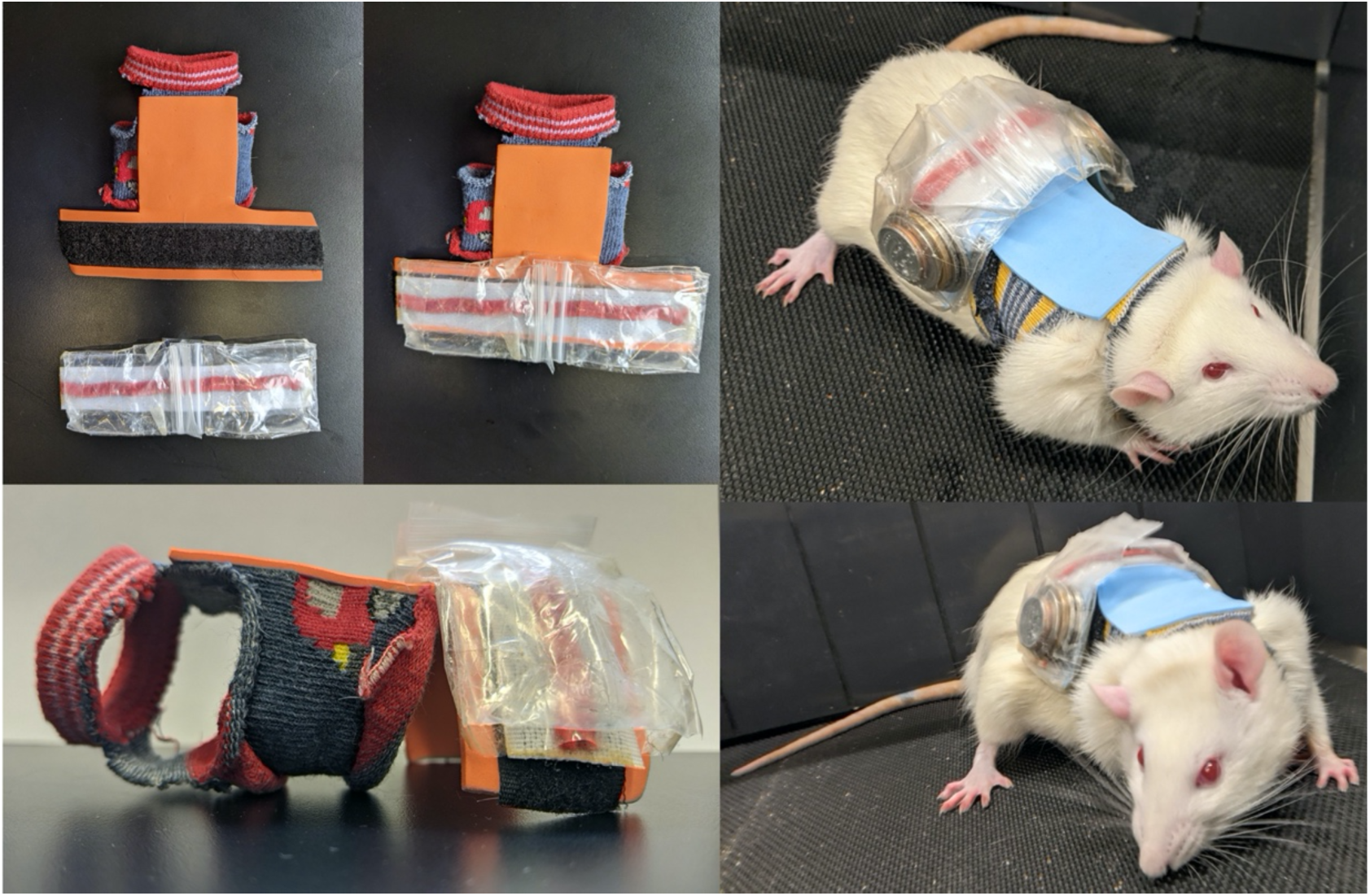
Rat weighted vest design (left) and a rat wearing a vest containing 15% of its body mass at week 4 of training (right).

### Training program

The training program was modelled after those by Butterfield et al. (2005) and Chen et al. (2020), who saw SSN adaptations following 2-4 weeks of downhill running training in the rat vastus intermedius and soleus, respectively, compared to control and uphill running groups. Two weeks prior to training, rats were handled for 1 hour on 3 consecutive days to reduce their stress levels when later applying the vests. The next week (i.e., 1 week before training), the rats underwent 5 consecutive days of familiarization sessions on the treadmill, each consisting of three 3-minute sets at a 0° grade and 10-12 m/min speed, with 2 minutes of rest between each set. In the first two familiarization sessions they did not wear vests, in the third session they wore an unweighted vest for 1/3 sets, in the fourth session they wore an unweighted vest for 2/3 sets, and in the fifth session they wore an unweighted vest for all 3 sets. Rats who were not compliant to the treadmill after 3 days of attempted familiarization were made controls instead.

Previous models of eccentric-biased resistance training in rats, rabbits, and mice optimized muscle architectural adaptations with lower compared to higher training frequencies (Butterfield and Herzog, 2006; Morais et al., 2020; Ochi et al., 2007; Wong and Booth, 1988; Zhu et al., 2021), so rats ran 3 days per week (Monday, Wednesday, and Friday). Rats ran on an EXER 3/6 animal treadmill (Columbus Instruments, Columbus, OH, USA) set to a 15° decline. Rats ran in 5-minute bouts, separated by 2 minutes of rest. They completed 3 bouts on the first day and increased by 2 bouts/day up to the 7-bout target (35 minutes total) on the third day of training, which persisted for the remainder of the training period. Rats began each training session at a speed of 10 m/min, which was increased by 1 m/min up to the 16 m/min target (Chen et al., 2020). Progressive loading was employed by adding weight equivalent to 5% of the rat’s body mass during the first week, 10% in the second week, 15% in the third week, and 15% readjusted to the new body mass in the fourth week. The training efficacy and safety of this approach was modelled after a weighted vest training review which reported added weights of 5%-20% body mass (Macadam et al., 2019), and work by Schilder et al.(2011), who added up to 36% of body mass to rats via backpacks to simulate obesity. 15% of body mass was the maximum weight used in the present study because during piloting we found that adding 20% of body mass was too cumbersome for the animals while running. Each training session took place at approximately the same time of day (between 9 and 11 AM).

### Experimental setup

Immediately following sacrifice, the soleus was carefully harvested from the right hindlimb (Ma and Irving, 2019). Silk-braided sutures (USP 2-0, metric 3) were tied along the musculotendinous junctions and mounted to the force transducer and length controller in the 806D Rat Apparatus (Aurora Scientific, Aurora, ON, Canada). The muscles were bathed in a ∼26°C Tyrode solution with a pH of ∼7.4 (121 mM NaCL, 24 mM NaHCO_3_, 5.5 mM D-Glucose, 5 mM KCl, 1.8 mM CaCl_2_, 0.5 mM MgCl_2_, 0.4 mM NaH_2_PO_4_, 0.1 mM EDTA) that was bubbled (Bonetta et al., 2015; Cheng and Westerblad, 2017) with a 95% O_2_/5% CO_2_ gas mixture (Praxair Canada Inc., Kitchener, ON, Canada). A 701C High-Powered, Bi-Phase Stimulator (Aurora Scientific, Aurora, ON, Canada) was used to evoke all contractions via two parallel platinum electrodes submerged in the solution, situated on either side of the muscle. Force, length, and stimulus trigger data were all sampled at 1000 Hz with a 605A Dynamic Muscle Data Acquisition and Analysis System (Aurora Scientific, Aurora, ON, Canada). All data were analyzed with the 615A Dynamic Muscle Control and Analysis High Throughput (DMC/DMA-HT) software suite (Aurora Scientific, Aurora, ON, Canada).

### Mechanical testing

Mechanical testing of the soleus proceeded from Protocol A to D. As a starting point for approximating optimal length, the soleus was passively set to a taut length that resulted in ∼0.075 N of resting tension prior to beginning any protocols (Chen et al., 2020).

*Protocol A: Twitch current.* Single 1.25-ms pulses were delivered in increasing increments of 0.5 mA (starting at 1 mA) until peak twitch force was elicited. The current for peak twitch force was used in the remainder of mechanical testing.

*Protocol B: Force-length relationships.* The muscle was lengthened in 1-mm increments, with maximal tetanic stimulation (0.3 ms pulse width, 1 s duration, 100 Hz frequency) delivered at each new length (Chen et al., 2020; Heslinga and Huijing, 1993). This was repeated until the muscle length that produced peak tetanic force (L_O_) was reached. Construction of an active force-length relationship was then completed to at least ±2 mm with respect to L_O_. To further approximate/confirm L_O_ for the subsequent work loops, the muscle was stimulated at the initial L_O_ ± 0.5 mm. The final length for L_O_ was measured from tie to tie (i.e., musculotendinous junction to musculotendinous junction) using 150 mm (0.01 mm resolution) analog calipers (Marathon, Vaughan, ON, Canada). Prior to each tetanic stimulation, 2 minutes (to allow for dissipation of force transients owing to stress-relaxation) of resting force were obtained for construction of a passive force-length relationship, and following each tetanic stimulation, the muscle was returned to the original taut length (Heslinga and Huijing, 1993).

*Protocol C: Active and passive work loops.* Sinusoidal muscle length changes were imposed about L_O_ for each of 1.5, 2, and 3-Hz cycle frequencies (i.e., the frequency of the length sinusoid) and 1, 3, 5, and 7-mm strains (e.g., a 3-mm strain indicates a displacement of ±1.5 mm from L_O_). After piloting, we decided to not test the 7-mm strain at the 3-Hz cycle frequency because the muscle sometimes slipped out of its ties with this combination of speed and stretch.

The range of cycle frequencies was chosen because they correspond to a wide range of treadmill running speeds for the rat (<13 to 67 m/min) and are consistent with cycle frequencies that occur during spontaneous running (maximum 4 Hz) (Caiozzo and Baldwin, 1997; Nicolopoulos-Stournaras and Iles, 1983; Roy et al., 1991). The range of strains was chosen because they fall within the maximum excursion of the rat soleus *in vivo* (∼10 mm) (Woittiez et al., 1985). The cycle frequency and strain ranges we used are also comparable to those used in previous rat soleus work loop studies (Caiozzo and Baldwin, 1997; Swoap et al., 1997).

A set of 3 consecutive work loops was performed at each cycle frequency: 1 passive work loop with no electrical stimulation to determine the work done against passive elements in the muscle (Cox et al., 2000; James et al., 1995); and 2 active work loops with stimulation (0.3 ms pulse width, 100 Hz frequency) set to begin at the onset of shortening (Swoap et al., 1997) and last for 70% of the shortening phase (i.e., duty cycle = 0.7). Pilot testing determined that 2 consecutive active work loops sufficiently determined optimal net work: in a set of 3 active work loops, the highest net work occurred in the 1^st^ or 2^nd^ loop more than 80% of the time, and net work did not differ between each loop by >10%. Employing only 2 active work loops per set was also desirable for minimizing the development of muscle fatigue and deterioration of tissue viability. Two minutes of rest were always provided before proceeding to the next set of work loops (Swoap et al., 1997).

*Protocol D: Assessment for fatigue-induced force loss.* Two minutes after the final work loop, a maximal tetanic contraction performed at L_O_, and a work loop at 1.5 Hz and 3 mm (i.e., the loop determined to usually produce maximal net work output during piloting) were performed to ensure that fatigue did not interfere with results. Fatigue was defined as a >10% drop in these parameters compared to baseline, and no muscles experienced this.

### Limitations in addressing concerns of tissue viability

The present study only characterized the force-length relationship up to 2 mm on either side of L_O_, at 1-mm intervals. This methodological limitation may have prevented us from observing adaptations such as a widening of the force-length relationship plateau region (Akagi et al., 2020) or widening of the whole force-length relationship (i.e., the operating range for active force) (Alder et al., 1958; De Koning et al., 1987; Woittiez et al., 1986). Such information may have been relevant in interpreting the work loop results, as we would be able to better elucidate whether trained muscles were acting at near-optimal force production across a wider range of muscle lengths than controls. However, due to concerns of tissue viability we wanted to minimize time spent on the force-length relationship prior to work loop protocols. We also only employed 1 passive work loop and 2 active work loops per set in effort to preserve tissue viability for the duration of the experiment. There is likely some viscosity present during the first loop that would be reduced in later cycles and yield different net work output values. However, previous work loop studies on the rat soleus also employed as few as 3 total work loops per set in assessing net work output across a range of strains and cycle frequencies (Swoap et al., 1997). Additionally, as our work loop methods were consistent across work loop conditions (i.e., cycle frequency, strain) and between groups, it is unlikely that they confounded comparisons across cycle frequencies, strains, and groups in this study.

### Muscle architecture and serial sarcomere number estimations

Following mechanical testing, the muscle was removed from the bath, weighed, then passively stretched to the L_O_ determined in Protocol B and tied to a wooden stick. The muscle was then fixed in 10% phosphate-buffered formalin for 48 hours, rinsed with phosphate-buffered saline, and digested in 30% nitric acid for 6-8 hours to remove connective tissue and allow individual muscle fascicles to be teased out (Butterfield et al., 2005; Chen et al., 2020). To obtain a global measure of SSN, 6 fascicles were obtained lateral to medial, superficial, and deep across the muscle. Dissected fascicles were placed on glass microslides (VWR International, USA), then fascicle lengths were measured using ImageJ software (version 1.53f, National Institutes of Health, USA) from pictures captured by a level, tripod-mounted digital camera, with measurements calibrated to a ruler that was level with the fascicles. Sarcomere length measurements were taken at six different locations along each fascicle via laser diffraction (Coherent, Santa Clara, CA, USA) with a 5-mW diode laser (25 μm beam diameter, 635 nm wavelength) and custom LabVIEW program (Version 2011, National Instruments, Austin, TX, USA) (Lieber et al., 1984), for a total of 36 sarcomere length measurements per muscle. Serial sarcomere numbers were calculated as:

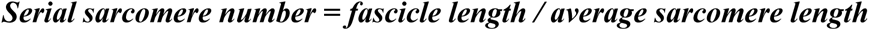

### Collagen content and solubility assay

The left soleus of 11 control and 12 trained rats was dissected and stored at -80°C until use in a hydroxyproline with collagen solubility assay, with pepsin-insoluble collagen quantifying the amount of collagen crosslinking (Brashear et al., 2020). The muscle was powdered in liquid nitrogen with a mortar and pestle, careful to remove any remaining tendon. The powdered muscle was then weighed and washed in 1 mL of phosphate-buffered saline and stirred for 30 minutes at 4°C. Non-crosslinked collagen was digested in a 1:10 mass (mg of powdered tissue):volume (µL) solution of 0.5 M acetic acid with 1 mg/mL pepsin, stirring overnight at 4°C. After centrifuging at 16000 g for 30 minutes at 4°C, the supernatant was collected as the pepsin-soluble fraction (non-crosslinked collagen), and the pellet was kept as the pepsin-insoluble fraction (crosslinked collagen). The two separate fractions were hydrolyzed overnight in 0.5 mL of 6 M hydrochloric acid at 105°C. 10 µL of hydrolysate were mixed with 150 µL of isopropanol followed by 75 µL of 1.4% chloramine-T (ThermoFisher) in citrate buffer (pH = 6.0) and oxidized at room temperature for 10 minutes. The samples were then mixed with 1 mL of a 3:13 solution of Ehrlich’s reagent (1.5 g of 4-[dimethylamino] benzaldehyde [ThermoFisher]; 5 mL ethanol; 337 µL sulfuric acid) to isopropanol and incubated for 30 min at 58°C. Quantification was determined by extinction measurement of the resulting solution at 550 nm. A standard curve (0-1000 µM trans-4-hydroxy-L-proline; Fisher) was included in each assay (average linear R^2^ = 0.98), and samples were run in triplicate (average CV = 0.098). Hydroxyproline concentrations in samples were determined by interpolation from the linear equation of the corresponding assay’s standard curve. Results are reported as µg of hydroxyproline per mg of powdered tissue mass.

### Data and statistical analyses

For construction of active force-length relationships, passive force values were subtracted from total force at each length to estimate active force production. Passive force was taken from a 500-ms window prior to tetanic stimulation, and total force was taken as the maximum force produced in the tetanic contraction. Average active and passive force-length relationships for each group were constructed by averaging force values in length categories with respect to L_O_ (e.g., L_O_ –2, –1, 0, +1, +2 mm).

From each work loop, Aurora Scientific software calculated work of shortening and work of lengthening (the integrals under the shortening and lengthening curves, respectively, in a force-length plot) and net work output of a whole cycle (the area inside the whole shortening and lengthening cycle; work of shortening + work of lengthening, with work in the shortening direction defined as positive). Of the two active work loops in a given set, the loop that produced the highest net work was chosen for statistical analyses (Swoap et al., 1997). Since a relatively long duty cycle (i.e., 0.7) may not have allowed full relaxation prior to lengthening in the active work loops, the net work from a work loop with a more optimal duty cycle was also estimated by adding the work of shortening (positive) in the active work loop and the work of lengthening (negative) in the corresponding passive work loop. These estimated values of optimal net work output will henceforth be referred to as the “estimated optimal net work output.”

All force and work values were normalized to the muscle’s physiological cross-sectional area (PSCA; cm^2^) to obtain specific values (expressed as units/cm^2^). PCSA was calculated as:

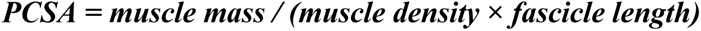

With muscle density assumed to be 1.112 g/cm^3^ (Ward and Lieber, 2005).

All statistical analyses were conducted in IBM SPSS statistics version 26. Two-tailed t-tests were used to compare body weights between control and trained rats throughout the training period (i.e., at ∼14, 15, 16, 17, and 18 weeks old). A one-way analysis of variance (ANOVA) with a Holm-Sidak correction for all pairwise comparisons was used to compare FL, SL, SSN, L_O_, muscle weight, PCSA, total collagen, and % insoluble collagen between trained and control rats. Since characterization of longitudinal muscle growth was the primary objective of this study, post-hoc power analyses (t-tests, difference between two independent means; G*Power software) were performed on FL, SL, and SSN, with 80% indicating statistical power.

One control rat was determined to be an outlier for L_O_ (being 2.09 mm greater than the third quartile + 1.5 times the interquartile range) and was removed from all force-length relationship analyses (new sample size: n = 16 control, n = 12 training). Two-way ANOVAs (group [training, control] × length with respect to L_O_ [L_O_ -2, -1, +0, +1, +2 mm]) with Holm-Sidak corrections were used to compare active and passive force across the force-length relationship, and between groups. Differences in active force at each length with respect to L_O_ were then compared using one-way ANOVAs with Holm-Sidak corrections. Since the force-length relationship was shifted rightward by on average ∼1 mm (see results), to determine if training reduced passive force at a given muscle length, passive force values of control rats at a length with respect to L_O_ were compared to passive force at one less than that length for trained rats (e.g., control passive force at L_O_ + 2 mm [∼26.7 mm] was compared to trained passive force at L_O_ + 1 mm [∼26.7 mm], control passive force at L_O_ + 1 mm [∼25.7 mm] was compared to trained passive force at L_O_ [∼25.7 mm], and so on) using a one-way ANOVA with a Holm-Sidak correction.

A three-way ANOVA (group [training, control] × cycle frequency [1.5, 2, 3 Hz] × strain [1, 3, 5, 7 mm]) with a Holm-Sidak correction for all pairwise comparisons was used to compare all work loop parameters from the passive and active work loops (work of shortening, work of lengthening, net work output, estimated optimal net work output). An effect of group defined a difference between control and trained rat work loops. Where effects of group were detected, one-way ANOVAs with Holm-Sidak corrections were employed to compare work loop parameters between control and trained rats at specific cycle frequencies/strains. Two-tailed paired t-tests were used to compare net work output between the recorded work loops and the estimated optimal work loops.

Regression analyses between net work output and SSN, FL, SL, and maximum isometric force normalized to PCSA in both groups combined were performed to elucidate which training-induced adaptations contributed to improvements in net work output. Multiple linear regression with the variables that showed significant relationships with net work output was also performed to further understand the relative importance of each in net work output.

Significance was set at *α* = 0.05. Effect sizes from two and three-way ANOVAs are reported as the partial eta squared (η_p_^2^), and from one-way ANOVAs as Cohen’s *d* (small effect = 0.2, medium effect = 0.5, large effect = 0.8) where significance was detected. Data are reported as mean ± standard deviation in text and mean ± standard error in figures.

## Results

### Body weight, muscle weight, and PCSA

As shown in Figure S1, there were no differences in body weight between control and trained rats at any time points. As shown in Figure 2A-B, there were also no differences in muscle wet weight (F(1,29) = 0.26, *P* = 0.62) or PCSA between trained and control rats (F(1,29) = 2.33, *P* = 0.14).

**Figure 2:**
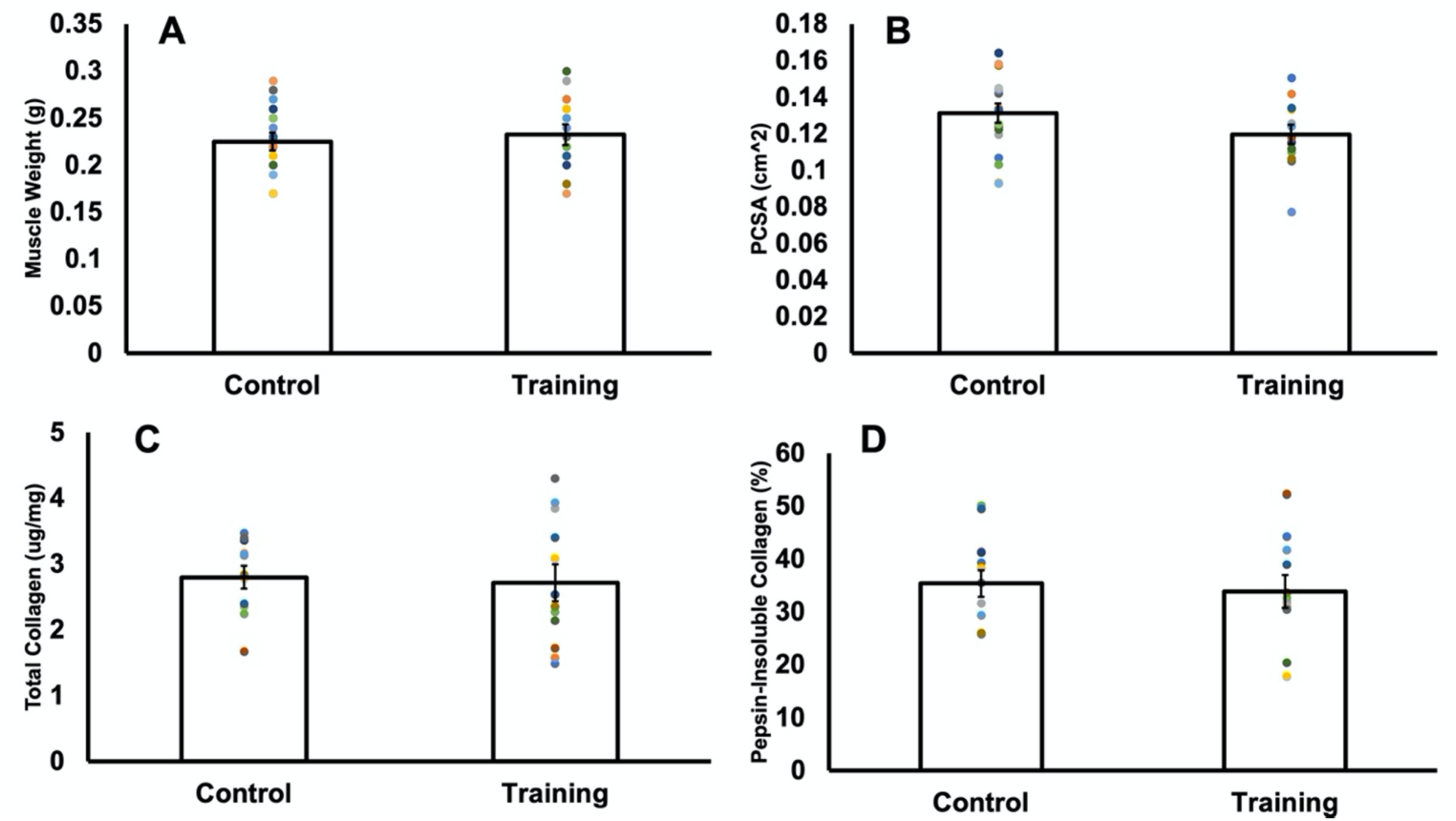
Comparison of muscle wet weight (A), physiological cross-sectional area (PCSA) (B), total collagen concentration (C), and the percentage represented by pepsin-insoluble collagen (D) in control versus trained rats. Data are reported as mean ± standard error (A-B: n = 18 control, n = 13 training; C-D: n = 11 control, n = 12 training). No significant differences (*P* > 0.05) were found between control and training for any of these variables.

### Collagen content and crosslinking

There were no differences in total hydroxyproline concentration (F(1,21) = 0.06, *P* = 0.81) (Figure 2C) or percent pepsin-insoluble collagen (F(1,21) = 0.14, *P* = 0.71) (Figure 2D) between trained and control rats, indicating no differences in collagen content or the amount of crosslinked collagen.

### Longitudinal muscle architecture

For average FL, 95% power was achieved with 18 control and 13 trained rats. FL was 13% longer (F(1,29) = 13.44, *P* < 0.01, *d* = 1.38) in trained (17.52 ± 1.60 mm, 95% CI [16.55,18.49]) than control rats (15.47 ± 1.49 mm, 95% CI [14.72,16.21]), implying a training-induced increase in FL (Figure 3A).

**Figure 3:**
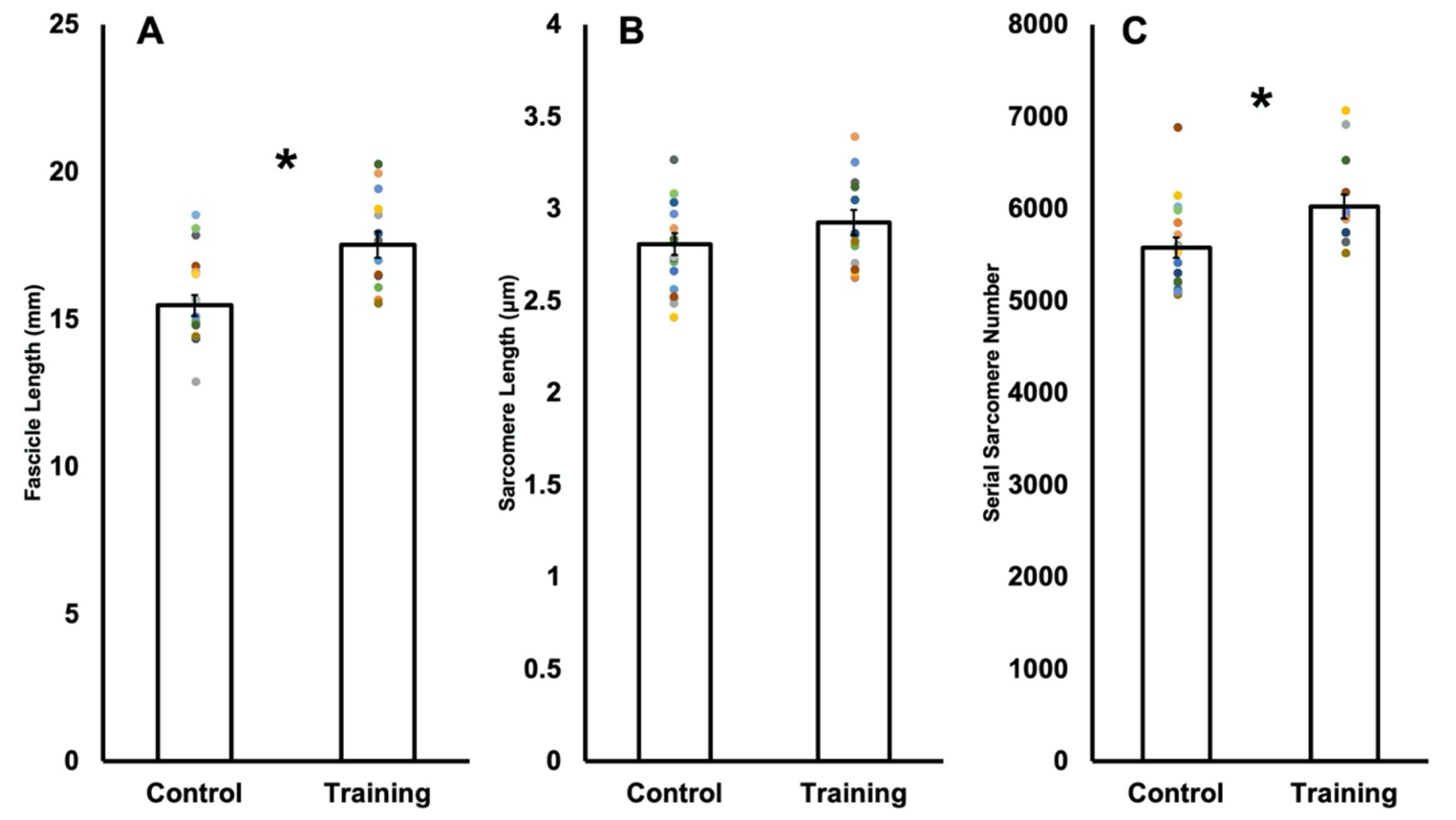
Comparison of fascicle length (A), sarcomere length (B), and serial sarcomere number (C) in control versus trained rats. Data are reported as mean ± standard error (n = 18 control, n = 13 training). *Significant difference (*P* < 0.05) between control and training.

For SL, 80% power was not achieved using average SL in 18 control and 13 trained rats (Power = 35%). With these sample sizes, SL did not differ (F(1,29) = 1.73, *P* = 0.20) between trained (2.93 ± 0.25 μm, 95% CI [2.78,3.07]) and control rats (2.81 ± 0.25 μm, 95% CI[2.69,2.94]) (Figure 3B). A follow-up a-priori power analysis indicated 67 control and 49 trained rats would be required to achieve statistical power for SL, which would not be feasible within time and ethical constraints of this research. To obtain data closer to this sample size, we treated each fascicle independently, amounting to 106 control and 78 trained measurements of average SL. Viewed this way, SL was 5% higher (F(1,182) = 8.75, *P* < 0.01, *d* = 0.44) in trained (2.94 ± 0.29 μm, 95% CI [2.87, 3.01]) than control rats (2.81 ± 0.29 μm, 95% CI [2.76, 2.87]) (Figure 4A). Therefore, there is evidence that training increased SL, however, only when using a larger sample size representative of individual fascicles as opposed to individual animals.

**Figure 4:**
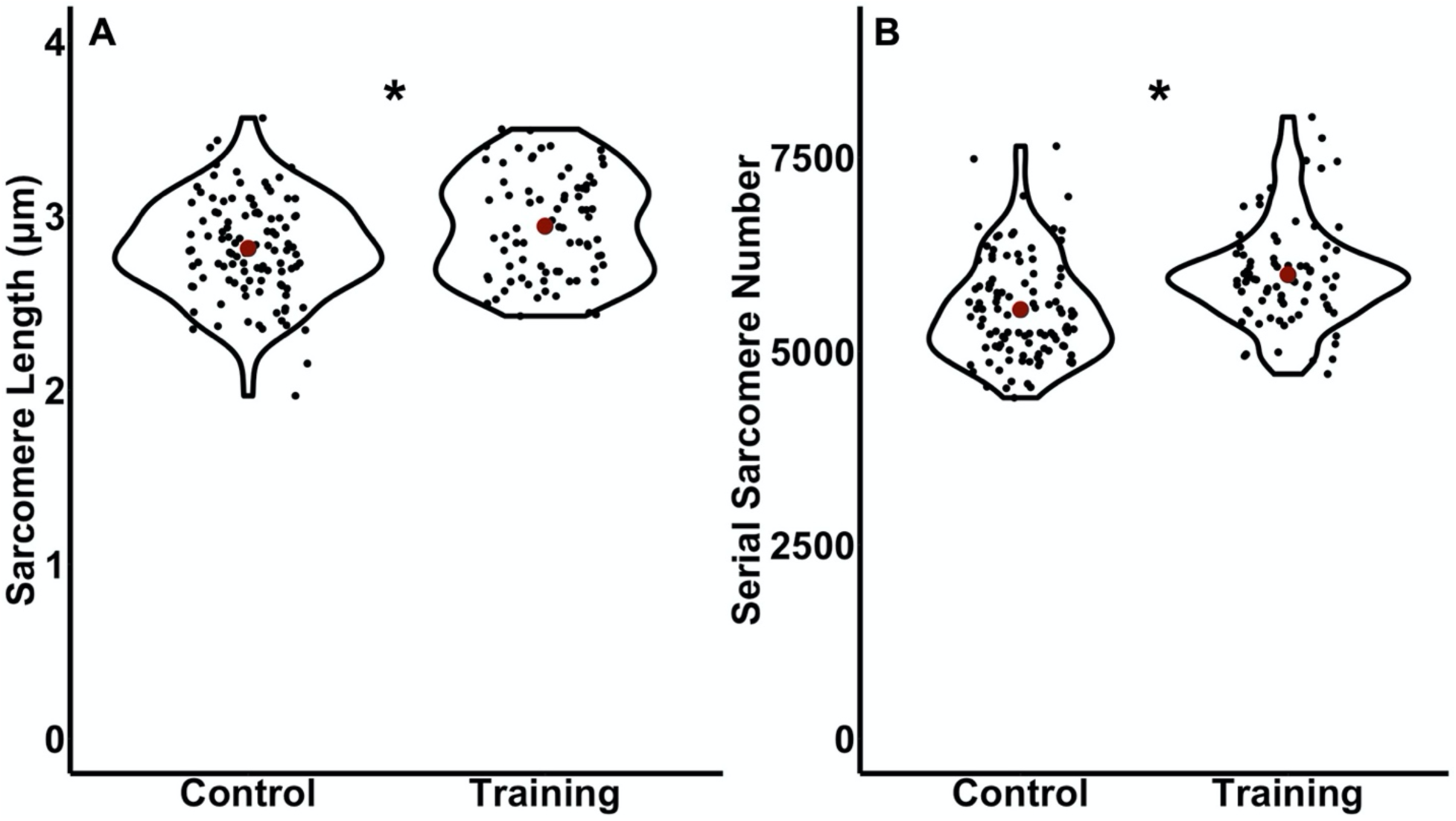
Violin plot comparisons of average sarcomere length (A) and serial sarcomere number (B) in control versus trained rats with the sample size adjusted to treat each fascicle independently. Red dots represent the mean. *Significant difference (*P* < 0.05) between control and training.

For average SSN, 82% power was achieved with 18 control and 13 trained rats. SSN was 8% greater (F(1,29) = 6.89, *P* = 0.01, *d* = 0.99) in trained (6024.45 ± 473.38, 95% CI [5738.39, 6310.51]) compared to control rats (5577.03 ± 464.52 mm, 95% CI[5346.03,5808.03]) (Figure 3C). Figure 4B also shows that SSN was 8% greater in trained compared to control rats when viewing the measurements from all fascicles (106 control and 78 trained fascicles) (F(1,182) = 21.26, *P* < 0.01, *d* = 0.69).

### Force-length relationships

While L_O_ was on average ∼1 mm longer in trained (25.72 ± 1.27 mm) than control rats (24.69 ± 2.70 mm) (Figure 5A and C), there was not a significant difference (F(1,26) = 1.48, *P* = 0.23). There was an effect of group on specific active force (F(1,135) = 27.09, *P* < 0.01, η_p_^2^ = 0.18), with one-way ANOVA showing specific active force was ∼50% greater in trained than control rats at L_O_ -1 mm (F(1,26) = 5.77, *P* = 0.02, *d* = 0.95), L_O_ (F(1,26) = 5.88, *P*= 0.02, *d* = 0.96), L_O_ +1 mm (F(1,26) = 6.19, *P* = 0.02, *d* = 0.99), and L_O_ +2 mm (F(1,22) = 6.15, *P* = 0.02, *d* = 1.07). There was no effect of muscle length with respect to L_O_ on specific active force (F(4,135) = 0.36, *P* = 0.84), suggesting active force did not significantly differ between L_O_ and the other muscle lengths tested, however, when comparing force values normalized to maximum force, there was an effect of muscle length with respect to L_O_ (F(4,133) = 26.37, *P* < 0.01, η_p_^2^ = 0.46), and force differed from force at adjacent points at all muscle lengths tested (all comparisons *P* < 0.01) (Figure 5A).

**Figure 5:**
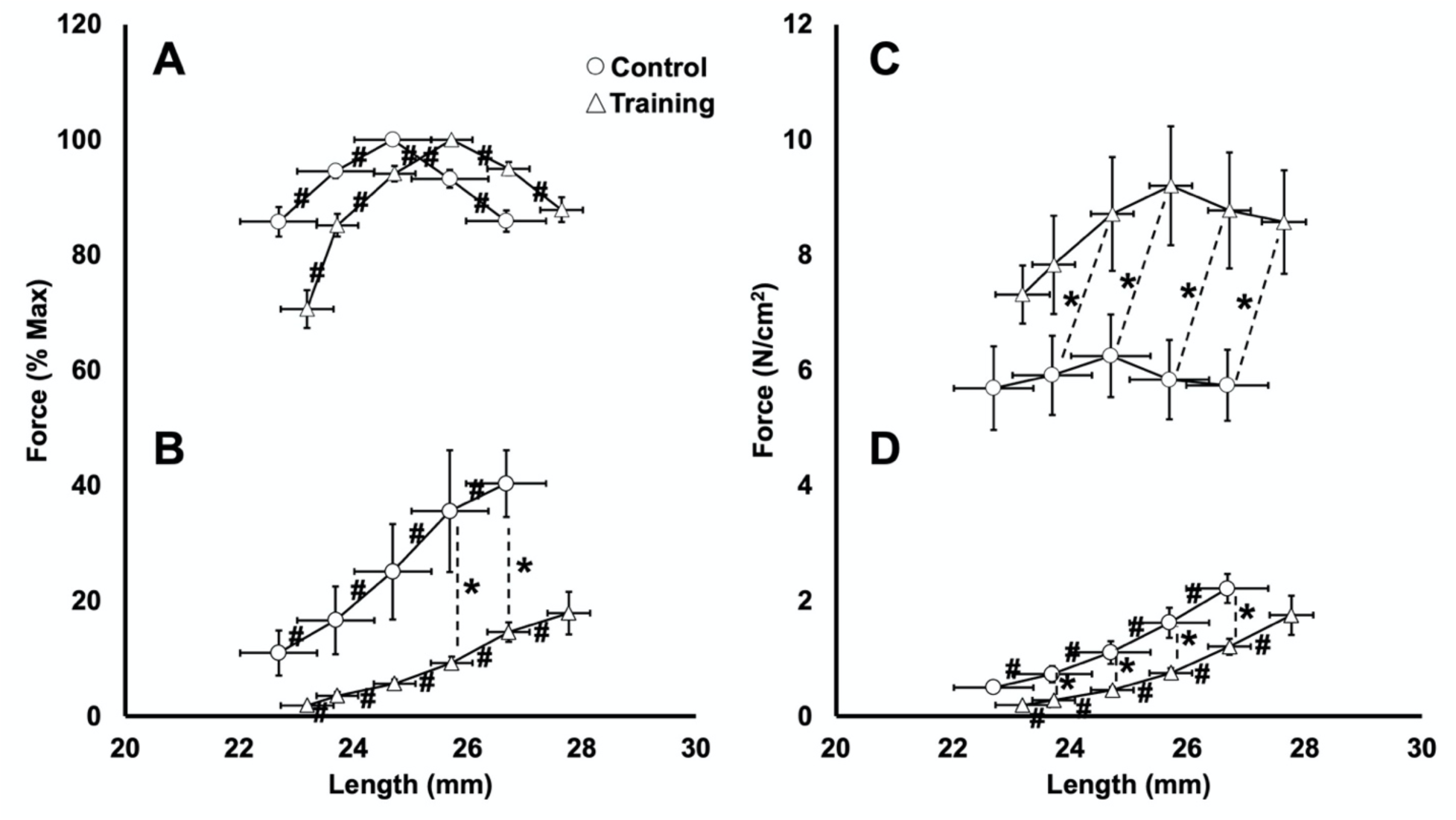
Comparison of average active (A and C) and passive (B and D) force-length relationships in control versus trained rats, expressed as percent of maximum active force (left) and force normalized to physiological cross-sectional area (right). Data are reported as mean ± standard error (n = 16 control, n = 12 training). *Significant difference (*P* < 0.05) between control and training. #Significant difference in force between muscle lengths.

As shown in Figure 5B and D, the average passive force-length relationship also shifted rightward in trained compared to control rats. There was an effect of group on both specific force (F(1,136) = 29.52, *P* < 0.01, η_p_^2^ = 0.22) and force normalized to maximum active force (F(1,133) = 12.85, *P* < 0.01, η_p_^2^ = 0.10) in the passive force-length relationship. Particularly, specific passive force was 45-62% less in trained than control rats at ∼23.7 mm (F(1,26) = 7.34, *P* = 0.01, *d* = 1.07), ∼24.7 mm (F(1,26) = 7.53, *P* = 0.01, *d* = 1.09), ∼25.7 mm (F(1,26) = 7.39, *P* = 0.01, *d* = 1.08), and ∼26.7 mm (F(1,25) = 8.63, *P* < 0.01, *d* = 1.16) (Figure 6D). Passive force normalized to maximum was 26% less in trained than control rats at ∼25.7 mm (F(1,26) = 4.61, *P* = 0.04, *d* = 0.85) and ∼26.7 mm (F(1,25) = 14.10, *P* < 0.01, *d* = 1.51) (Figure 6B). Last, there was an effect of muscle length with respect to L_O_ on specific passive force (F(4,135) = 26.10, *P* < 0.01, η_p_^2^ = 0.46) and passive force normalized to maximum (F(4,133) = 4.41, *P* < 0.01, η_p_^2^ = 0.13), such that force differed between adjacent points at all muscle lengths tested (all comparisons *P* < 0.05), confirming passive force increased with increasing muscle length.

**Figure 6:**
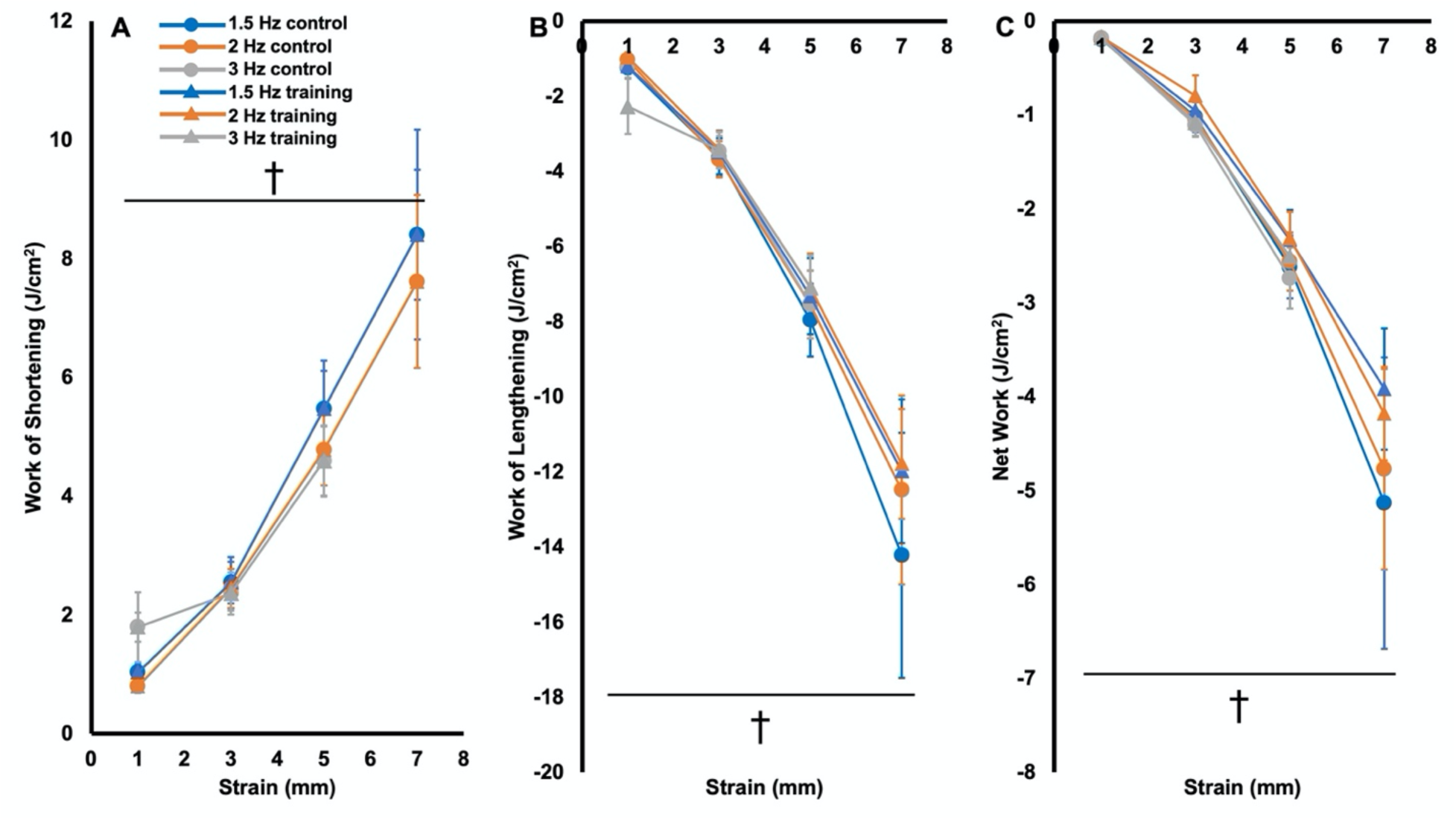
Work of shortening (A), work of lengthening (B), and net work output (C) of the passive (i.e., no stimulation) work loops in trained and control rats. Data are reported as mean ± standard error (n = 16 control, n = 12 training). †Significant difference across strains.

### Passive work loops

All passive work loops were clockwise in shape such that work of lengthening exceeded work of shortening (Figure S2). There were no effects of group (F(1,304) = 0.00-2.07, *P* = 0.15-0.99), nor group × cycle frequency × strain (F(5,304) = 0.08-0.10, *P* = 0.99-1.00), group × cycle frequency (F(2,304) = 0.04-0.14, *P* = 0.87-0.96), or group × strain (F(3,304) = 0.09-0.67, *P* = 0.57-0.97) interactions for work of shortening, work of lengthening, or net work output, indicating similar passive work loops between control and trained rats at each cycle frequency and strain (Figure 6).

There were no effects of cycle frequency (F(2,304) = 0.11-0.73, *P* = 0.48-0.90) nor cycle frequency × strain interactions (F(5,304) = 0.02-0.38, *P* = 0.86-0.98) on work of shortening, work of lengthening, or net work output in the passive work loops, indicating they did not differ between cycle frequencies at any strain. There were, however, effects of strain on all these parameters (F(3,304) = 71.10-85.57, *P* < 0.001, η_p_^2^ = 0.43-0.48), such that work of shortening and work of lengthening increased, and net work output decreased (i.e., became more negative) as strain increased (all comparisons *P* < 0.02) (Figure 6).

### Active work loops

Figure 7 shows representative work loop traces from 1 control and 1 trained rat. In general, work loops were mostly counterclockwise (i.e., containing primarily positive net work) up to work loops of at most 2 Hz and 3 mm, after which work loops became increasingly clockwise (i.e., containing negative work) with the work to re-lengthen the muscle exceeding the work of shortening.

**Figure 7:**
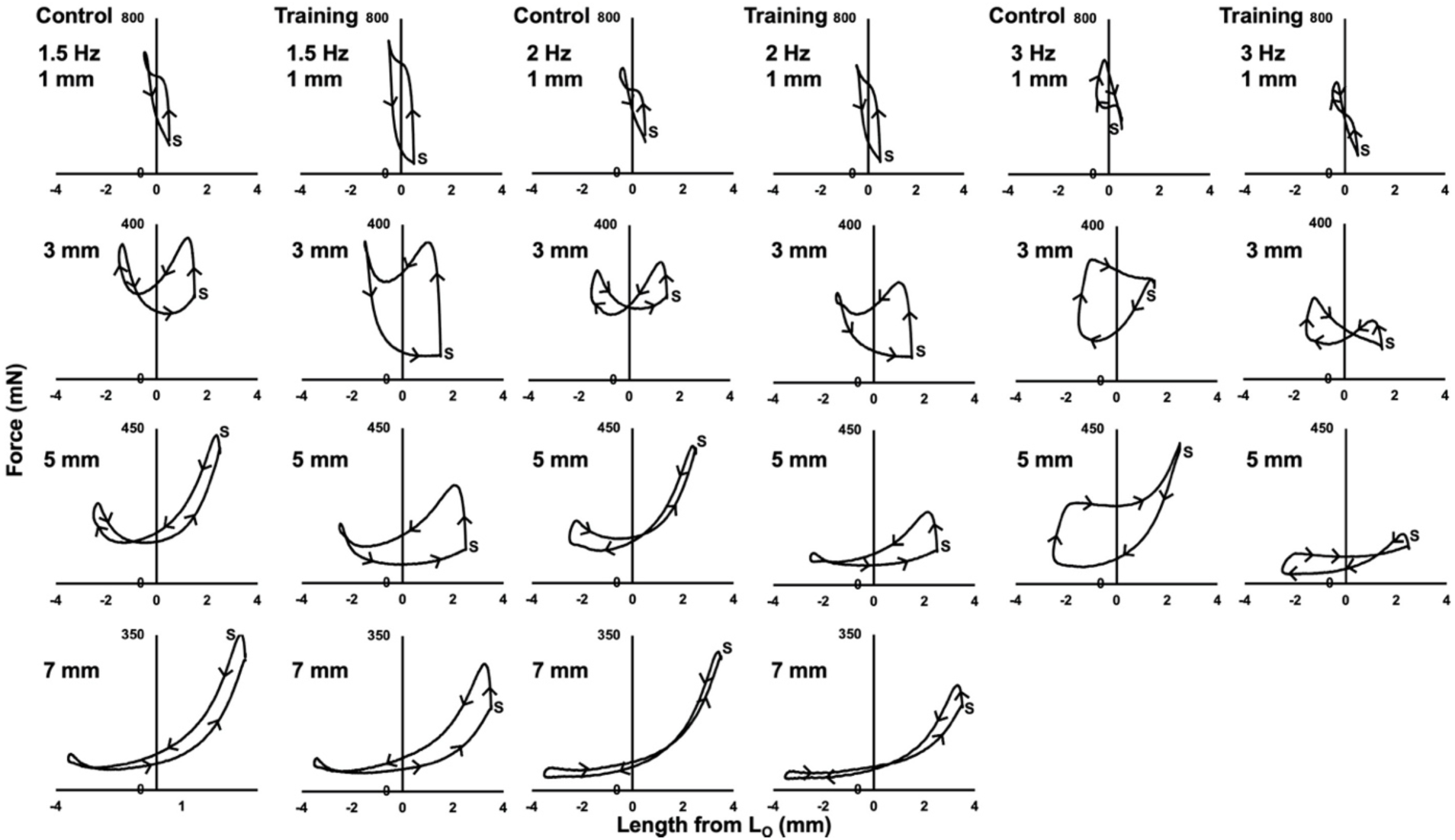
Representative active (i.e., stimulation during shortening) work loop traces from 1 control and 1 trained rat. *S* indicates the start of the cycle. Arrows indicate the direction of the cycle, with clockwise segments containing negative work and counterclockwise segments containing positive work.

As shown in Figure 8C, control rats on average produced maximal net work in the 1.5-Hz cycle at a 1-mm strain while trained rats produced maximal net work at 1.5 Hz and 3 mm, demonstrating a training-induced shift in the optimal strain at 1.5 Hz. Supporting training-induced changes in maximal net work output, there was an effect of group on net work output (F(1,304) = 9.19, *P* < 0.01, η_p_^2^ = 0.03), with trained rats producing on average 0.98 J/cm^2^ (95% CI [0.36, 1.61]) more net work than controls across all work loop conditions. However, there were no interactions of group × cycle frequency × strain (F(5,304) = 0.08 *P* = 1.00), group × cycle frequency (F(2,304) = 0.98, *P* = 0.38), or group × strain (F(3,304) = 0.47, *P* = 0.70), indicating the effect of group on net work output did not differ depending on cycle frequency and strain. Post-hoc tests showed the effect of group was most pronounced in 1.5-Hz work loops, with trained rats producing 101% greater net work output than controls at the 1-mm strain (F(1,26) = 6.42, *P* = 0.02, *d* = 1.00), 246% greater net work output at the 3-mm strain (F(1,26 = 4.92, *P* = 0.04, *d* = 0.88), and 424% greater net work output at the 5-mm strain (F(1,26 = 4.32, *P* < 0.05, *d* = 0.82) (Figure 8C).

**Figure 8:**
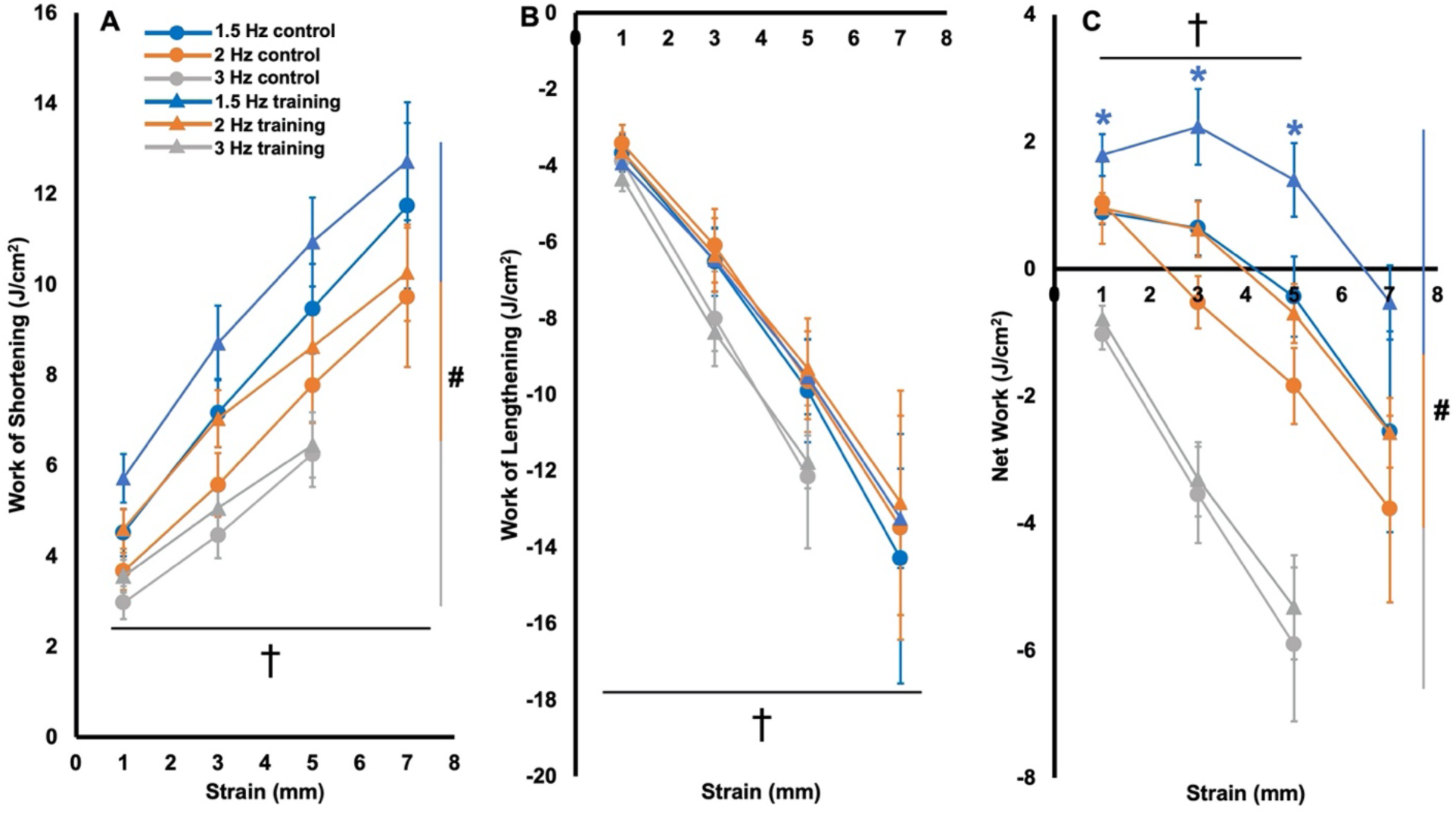
Work of shortening (A), work of lengthening (B), and net work output (C) of the active (i.e., stimulation during shortening) work loops in trained and control rats. Data are reported as mean ± standard error (n = 16 control, n = 12 training). *Significant difference (*P* < 0.05) between control and training at the colour-coded cycle frequency. #Significant difference across cycle frequencies. †Significant difference across strains.

There was an effect of group on work of shortening (F(1,304) = 5.52, *P* = 0.02, η_p_^2^ = 0.02) (Figure 8A) but not work of lengthening (F(1,304) = 0.06 *P* = 0.81) in the active work loops (Figure 8B), suggesting differences in work of shortening between groups contributed more to the differences in net work output, with trained rats producing on average 0.94 J/cm^2^ (95% CI [0.21, 1.68]) more work of shortening than controls across all work loops. Neither work of shortening or work of lengthening showed interactions of group × cycle frequency × strain (F(5,304) = 0.01-0.02 *P* = 1.00), group × cycle frequency (F(2,304) = 0.01-0.49, *P* = 0.61-0.99), or group × strain (F(3,304) = 0.12-0.14, *P* = 0.93-0.95). Despite there being an effect of group on work of shortening, post-hoc tests did not reveal significant differences in work of shortening between trained and control rats in any work loops.

There were also effects of cycle frequency on work of shortening and net work output, (F(2,304) = 20.37-59.51, *P* < 0.01, η_p_^2^ = 0.13-0.30), but not work of lengthening (F(2,304) = 2.42, *P* = 0.09). There were effects of strain on all these variables (F(3,304) = 26.95-43.68, *P* < 0.01, η_p_^2^ = 0.22-0.32). Specifically, work of shortening decreased with increasing cycle frequency (all comparisons *P <* 0.01), and work of shortening and work of lengthening both increased with strain (all comparisons *P* < 0.01) (Figure 8A-B). Net work output decreased with increasing cycle frequency (all comparisons *P* < 0.01), and with increasing strain past the optimal strain, except for between 5 mm and 7 mm (*P =* 1.00; all other comparisons *P* < 0.01 to 0.05) (Figure 8C).

To summarize all the above work loop data, there was an effect of group on net work output of the active work loops, with differences between control and trained rats most pronounced in the 1.5-Hz work loops at 1, 3, and 5-mm strains. Work of shortening and lengthening increased with strain in active and passive work loops, and net work output decreased (i.e. became more negative) with strain on either side of the optimal strain. Increasing cycle frequency decreased work of shortening and net work output, but did not change work of lengthening in the active work loops, and had no effect on any variables in the passive work loops.

### Relationships between net work output and serial sarcomere number, fascicle length, sarcomere length, and specific maximum isometric force

There were no significant relationships between SSN and net work output in any work loops. There were, however, significant relationships between net work output and FL (F(1,27) = 4.39-9.97, *P* = 0.01-0.04), and net work output and SL (F(1,27) = 4.87-13.77, *P* > 0.01-0.04) in the work loops that significantly differed between groups (i.e., 1.5 Hz at 1, 3, and 5-mm strains), such that FL explained 14% to 28% and SL explained 16% to 35% of the variation in net work output (Figure 9A-B). Regression analyses for all other work loops showed no significant relationships with FL or SL.

**Figure 9:**
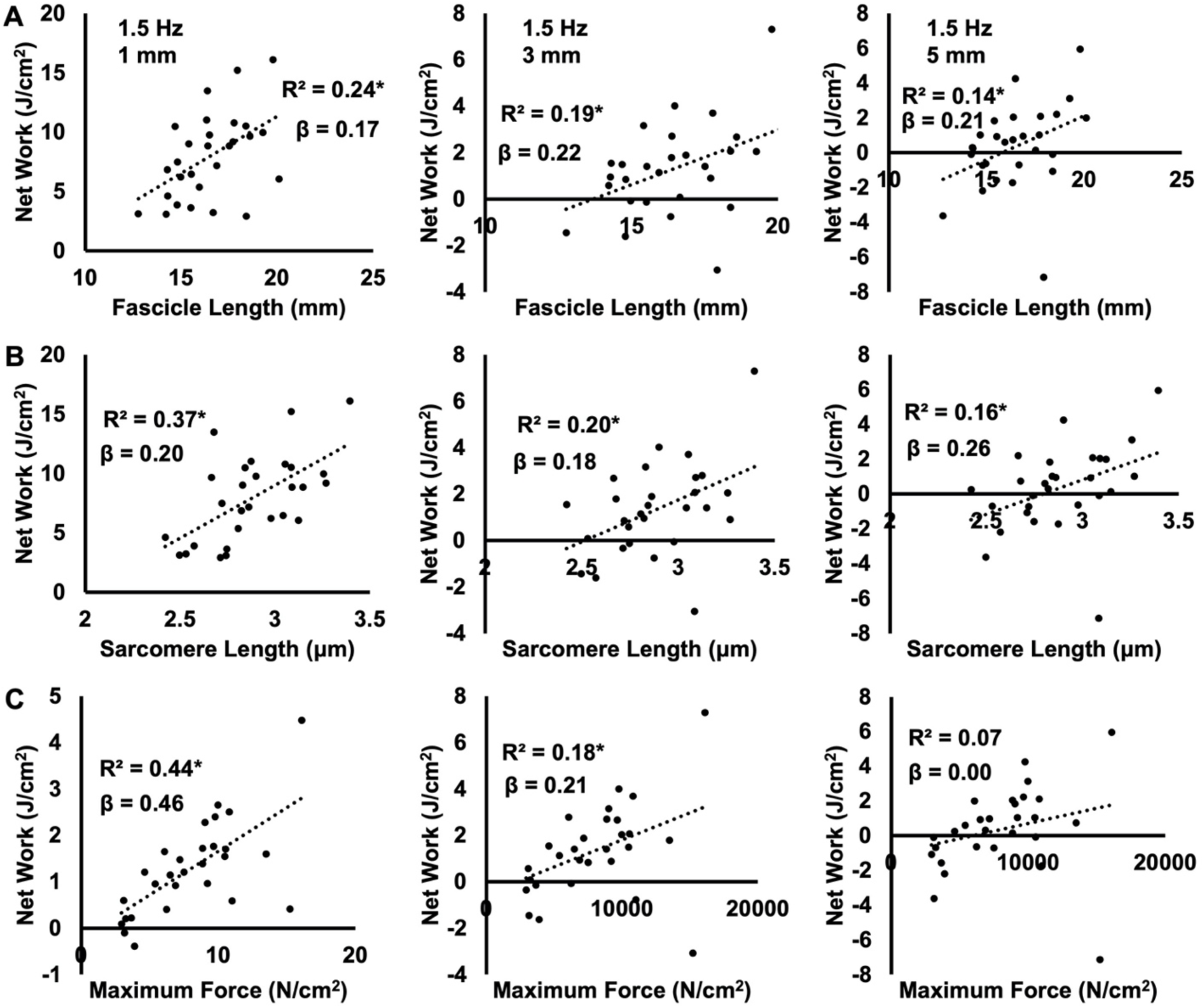
Plots of the relationships between net work output and fascicle length (A), sarcomere length (B), and maximum isometric force normalized to physiological cross-sectional area (C) in the 1.5-Hz work loops at strains of 1, 3, and 5 mm. *Significant relationship (*P* < 0.05).

There were significant relationships between specific maximum isometric force and net work output only in the 1.5-Hz work loops at 1 and 3-mm strains (F(1,27) = 5.59-20.55, *P* < 0.01-0.03), such that specific maximum isometric force explained 18-44% of the variation in net work output (Figure 9C). Beta coefficients from multiple linear regression revealed specific maximum isometric force contributed more to work loop performance at the 1-mm strain, while FL and SL were more important at the 3 and 5-mm strains (Figure 9).

## Discussion

This study assessed rat soleus architecture following 4 weeks of weighted downhill running training and aimed to relate muscle architectural adaptations to changes in dynamic contractile function, namely work loop performance. Our hypothesis that training would increase SSN, not change collagen content and crosslinking, and increase net work output especially in work loops with longer strains and faster cycle frequencies was partly correct. Comparing trained and control rats, FL and SSN increased with training, as did SL but only when treating every fascicle independently to increase the sample size. There were no differences in collagen parameters between groups, and improvements in net work output were most pronounced in 1.5-Hz work loops at strains of 1, 3, and 5 mm.

### Did weighted downhill running training induce sarcomerogenesis?

The FL, SL, and SSN values in this study are within ranges of those recorded previously for the rat soleus (Aoki et al., 2009; Baker and Hall-Craggs, 1978; Chen et al., 2020; Jakubiec-Puka and Carraro, 1991; Koh and Tidball, 1999). It should be noted that we only characterized global SSN of the soleus, thus any regional differences in SSN adaptations (Butterfield and Herzog, 2006) are beyond the scope of this study.

We observed an 8% training-induced increase in SSN. This magnitude of sarcomerogenesis is greater than what Chen et al. (2020) observed in the rat soleus following unweighted downhill running compared to controls (an insignificant +3%). Butterfield and Herzog (2006) showed the magnitude of SSN increase is strongly related to peak force developed during eccentric contractions. Therefore, compared to Chen et al. (2020), the weighted vests employed in the present study may have enhanced the eccentric load while running downhill, resulting in a greater stimulus for sarcomerogenesis. In the present study, rats also ran 3 days/week (Monday, Wednesday, Friday) rather than the 5 days/week (Monday-Friday) used by Chen et al. (2020), and the extra recovery days may have allowed more time for adaptations to occur between exercise bouts (Hyldahl and Hubal, 2014).

The most accepted hypothesis for sarcomerogenesis (i.e., an increase in SSN) stems from the active force-length relationship of muscle, whereby active force production is suboptimal in over-stretched positions due to limited actin-myosin binding (Gordon et al., 1966a; Gordon et al., 1966b), and observations that a muscle’s average SL operating range favours force production for particular task demands (Lieber and Ward, 2011; Lutz and Rome, 1994; Rome, 1994). It follows that if a muscle is repeatedly forced to operate with overstretched sarcomeres (i.e., with eccentric training), SSN would increase to maintain optimal actin-myosin overlap regions in that position (Koh, 1995; Lynn et al., 1998; Williams and Goldspink, 1978). These adaptations relate to a muscle’s ability to sense a change in tension (in this case increased tension at a long muscle length), then convert that mechanical signal into biochemical events regulating gene expression and protein synthesis (Aoki et al., 2009; Bogomolovas et al., 2021; Franchi et al., 2018; Herring et al., 1984; Soares et al., 2007; Spletter et al., 2018).

The 13% FL increase we observed with weighted downhill running training appears to have been driven by increases in both SSN (+8%) and SL (+4-5%). This may be interpreted in two ways. First, as optimal SL is generally believed to be constant within a species (Gokhin et al., 2014; Walker and Schrodt, 1974), the roles of increased SL and SSN in increasing FL are likely not mutually exclusive, but rather depend on the time course of adaptations (Herzog and Fontana, 2021; Zöllner et al., 2012). This perspective is illustrated by Barnett et al. (1980), who stretched the patagialis muscle of chickens for 10 days. At 24 hours, the researchers observed a 40% increase in biceps brachii SL, however, SL decreased back to normal by 72 hours, with instead an increase in SSN. Jahromi and Charlton (1979) explained this phenomenon by transverse myofibril splitting, whereby an overstretched sarcomere eventually splits at its H-zone, then two new sarcomeres develop from the split halves. It is therefore possible in the present study that at the 4- week mark, some sarcomeres were at the point of stretching (i.e., increasing SL), while others had separated such that new sarcomeres could be added (i.e., increasing SSN).

On the other hand, as we measured SL at L_O_, we may have observed an increase in optimal SL, which could imply elongation of the myofilaments within the sarcomere (Gokhin et al., 2014). In this regard, it is difficult to compare to previous studies on sarcomerogenesis, as most fixed their muscles in formalin at a set joint angle rather than at L_O_ (Roy et al., 1982; Salzano et al., 2018; Tabary et al., 1972; Williams and Goldspink, 1978). These studies reported on average approximately the same or shorter SLs in the experimental compared to control muscles. Therefore, it is hypothetically possible that, had they fixed the muscles at L_O_ (i.e., a longer length compared to the control muscle), they would have observed a longer optimal SL. This perspective is somewhat supported by other studies that fixed muscles at L_O_. Shrager et al. (2002) induced emphysema in the lungs of rats, then 5 months later performed lung volume reduction surgery to treat the emphysema. Another 5 months later, the diaphragms of rats who received lung volume reduction surgery had a 14% higher SSN and a 2% longer optimal SL than untreated rats (2.95 µm versus 2.88 µm). Coutinho et al. (2004) fixed the rat soleus at resting length, which they assumed to be L_O_, following 3 weeks of immobilization in a shortened position with intermittent stretching. Although they observed a decrease in SSN (due to being immobilized in a shortened position), they observed a ∼2% increase in SL compared to controls (2.2 µm versus 1.9 µm). Lastly, Chen et al. (2020) fixed the rat soleus at a measured L_O_ and observed on average a greater SL in downhill running rats than controls (2.77 ± 0.09 µm versus 2.75 ± 0.11 µm). Future research might aim to assess the time course of SL and SSN adaptations during training, or single-fiber testing to confirm optimal SLs to better understand these observations.

### Did weighted downhill running training induce muscle hypertrophy?

The rat soleus wet weight and PCSA values we observed are within the range of previous reports (Honda et al., 2018; Roy et al., 2002; Widrick et al., 2008). Increases in muscle wet weight and PCSA are believed to reflect muscle hypertrophy, and have been observed following mechanical loading in rats of similar age to those in the present study (Ochi et al., 2007; Roy et al., 1982; Wong and Booth, 1988). We, however, observed no difference in soleus wet weight or PCSA between trained and control rats, possibly because there was still a strong endurance component to our training program. Indeed, while the present study aimed to design a more resistance training-style program via the addition of weighted vests, rats still ran for a total of 35 minutes/session. Endurance training has been shown in humans to not alter anatomical CSA, which may provide insight into PCSA (Farup et al., 2012). If attempting to employ resistance training in rodents, future research might choose to investigate training programs closer to that of Zhu et al. (2021), in which mice pulled progressively overloaded carts down a track to failure, inducing large increases in muscle wet weight.

Muscle PCSA is believed to reflect parallel sarcomere number, and thus be proportional to maximum isometric force production (Haun et al., 2019; Narici et al., 2016). We observed ∼50% greater maximum isometric force normalized to PCSA at L_O_ and 3 other muscle lengths in trained compared to control rats. These data imply training improved characteristics associated with the intrinsic force-generating capacity of the muscle, such as calcium handling, force produced per crossbridge, or myosin density (i.e., number of crossbridges), which may improve with physical activity (Fitts et al., 1991; Xu et al., 2018) and resistance training (Łochyński et al., 2021).

### Did sarcomerogenesis impact force-length relations?

Although statistically insignificant, we observed on average a 4% (∼1 mm) greater L_O_ in trained than control rats. With an increase in SSN, L_O_ is expected to shift to a muscle length previously on the descending limb (Davis et al., 2020; Koh, 1995). Rightward shifts in L_O_ of 5-12% have been observed alongside 7-20% increases in SSN in studies on animals via immobilization in a stretched position, high-acceleration training, isokinetic eccentric training, and downhill running training (Butterfield and Herzog, 2006; Lynn et al., 1998; Salzano et al., 2018; Williams and Goldspink, 1978). The downhill running study (Lynn et al., 1998) assessed the joint torque-angle (i.e., not muscle force-length) relationship of the knee extensors, so is difficult to compare to the present study. Chen et al. (2020), though, employed downhill running training and observed no difference in rat soleus L_O_ compared to controls, and on average a 1.5% (also insignificant) greater L_O_ compared to uphill running rats. Like with SSN, the present study’s addition of weighted vests and extra rest days between training sessions may have amounted to a greater shift in rat soleus L_O_ compared to unweighted downhill running training.

Changes to the passive force-length relationship were more noticeable than changes to the active force-length relationship. Specifically, trained rats generated 45-62% less passive force at ∼23.7, 24.7, 25.7, and 26.7-mm muscle lengths. With a greater SSN, individual sarcomeres may stretch less during muscle displacement, leading to less passive force generated by sarcomeric proteins such as titin (Herbert and Gandevia, 2019). Noonan et al. (2020) demonstrated this with lower passive stress and passive elastic modulus at longer SLs in soleus single fibers of rats that ran downhill compared to uphill. Noonan et al. (2020) used the same rats as Chen et al. (2020), therefore the lower passive stress and elastic modulus appeared to be due to a 6% greater SSN.

The influence of sarcomerogenesis on the passive length-tension relationship is not always predictable due to the concurrent involvement of intramuscular collagen in passive force generation (Gillies and Lieber, 2011). For example, in studies on animals with sarcomerogenesis induced via surgically stretching a muscle, the passive force-length relationship steepens compared to controls due to concurrent increases in collagen content (Herbert and Balnave, 1993; Takahashi et al., 2012). Age-related increases in passive muscles stiffness are also related to increased collagen crosslinking in mice (Brashear et al., 2020). We observed no differences in collagen content or crosslinking in trained compared to control rats, with our values of hydroxyproline concentration falling into ranges of previous studies for the rat soleus (Karpakka et al., 1992; Sugama et al., 1999). Han et al. (1999) observed no change in collagen content in the rat soleus following downhill running, but also observed increases in some mRNAs and enzymes related to collagen synthesis; however, these were following only a single bout of exercise. Perhaps more comparable to the present study, Zimmerman et al. (1993) observed no changes in soleus collagen content or crosslinking in 20-week-old rats following 10 weeks of uphill running training. A rightward shift in the passive force-length relationship alongside increased SSN and no change in collagen content was also observed following passive stretch training of the rabbit soleus (De Jaeger et al., 2015). Altogether, the present study’s training-induced sarcomerogenesis appeared to reduce passive tension at longer muscle lengths.

### Is the work loop data comparable to previous studies on the rat soleus?

The present study’s active work loops employed stimulation duty cycles of 0.7, which is longer than optimal for all the cycle frequencies used (optimal = ∼0.6 at 1.5 Hz, ∼0.55 at 2 Hz, and ∼0.45 at 3 Hz (Swoap et al. 1997)). Consequently, full relaxation did not occur prior the lengthening phase in many of the active work loops, leading to some active lengthening force development and partly, if not fully (i.e., at 3 Hz), clockwise work loops (Figure 7). While we did not collect the most optimal possible work loops, stimulation parameters were consistent between control and trained rats, and as force was measured during muscle displacement, the data can be interpreted in the context of muscle architecture to address our research question. Furthermore, to bolster comparisons with prior studies, we estimated what net work output might have been under better stimulation conditions by subtracting work of lengthening in the passive work loops from work of shortening in the active work loops (Figure S3). When graphing these values of estimated optimal net work output across strain and cycle frequency, the graph is similar in shape to previous reports for the rat soleus (Caiozzo and Baldwin, 1997; Swoap et al., 1997). Additionally, the work loops we obtained that were counter-clockwise (e.g., Figure 7: Training, 2 Hz, 5 mm) were similar in shape to previous representative traces for the rat soleus (Swoap et al., 1997).

Net work output decreased with increasing cycle frequency and on either side of the optimal strain. At shorter strains, there is mathematically lower mechanical work, as work is the product of force and length change. At longer strains, the muscle undergoes greater stretch, thereby increasing passive force development in addition to active force development, and bringing the work of lengthening closer to or above work of shortening (Josephson, 1999; Swoap et al., 1997). Since work of shortening decreased with increasing cycle frequency while work of lengthening did not change, the decreased net work output with increasing cycle frequency can be attributed to the force-velocity relationship (i.e., force decreases with increasing shortening velocity due to less opportunity for crossbridge formation (Alcazar et al., 2019; Josephson, 1999)). Shorter muscle strain allows greater work at a faster cycle frequency because the shortening velocity (length change / time) is mathematically slower (James et al., 1995; Josephson and Stokes, 1989). The ability to perform at faster cycle frequencies can also depend on sarcoplasmic reticulum volume density (Lindstedt et al., 1998; Schaeffer and Lindstedt, 2013), and muscle fiber type (Swoap et al., 1997). Since the rat soleus is primarily a slow-twitch muscle (Armstrong and Phelps, 1984), it is understandable that net work output decreased considerably past a cycle frequency as slow as 1.5 Hz; this is consistent with previous work loops for the rat soleus (Caiozzo and Baldwin, 1997; Swoap et al., 1997).

One notable difference from Swoap et al. (1997) is they determined optimal strain for the rat soleus at 1.5 Hz to be 5 mm, while in the present study it was 1 to 3 mm. This may be because Swoap et al. (1997) tied the muscle at the distal end of the Achilles tendon, while we tied at the musculotendinous junctions. Incorporating tendon into work loops can improve net work output by adding forces generated by elastic recoil (Roberts, 2016), allowing muscle fibers to shorten at closer to optimal speeds (Lichtwark and Barclay, 2010), and buffering muscle damage during stretch (Griffiths, 1991). Swoap et al. (1997) also used a higher temperature (30°C versus 26°C), which can enhance force production in higher velocity/length change combinations (Ranatunga, 2018; Roots and Ranatunga, 2008). Altogether, while not perfect, our work loops still provide insight into how the rat soleus functions during cyclic contractions of various speeds and strains.

### Did sarcomerogenesis relate to improvements in work loop performance?

Our hypothesis that a longer FL and greater SSN induced by weighted downhill running would improve work loop performance, particularly at faster cycle frequencies and longer strains, was partly correct. An effect of group showed that trained rats produced overall greater net work output in the active work loops, with the most pronounced increases occurring in the 1.5-Hz loops at 1, 3, and 5 mm (Figure 8C). These improvements in net work seemed to be due to increases in work of shortening (i.e., active force during shortening) rather than decreases in work of lengthening (Figure 8A and B). Our hypothesis may only stand if optimal SL remained the same between control and trained rats. With increased SSN but the same SL, sarcomeres would stay in closer range to optimal SL, and shortening velocity of individual sarcomeres would be reduced for a given muscle excursion, improving active force production and thereby the mechanical work (Baxter et al., 2018; Drazan et al., 2019). We observed a 4-5% increase in optimal SL with training, so it is possible the sarcomere shortening velocities and operating ranges with respect to optimal SL did not change.

As shown in Table 1, we estimated the sarcomere operating ranges and shortening velocities in the 1.5-Hz work loops at strains of 1, 3, and 5 mm in trained and control rats. To do this, we first determined the fraction of the measured FLs compared to the whole-muscle L_O_’s (trained = 0.68, control = 0.63). By doing this, we can estimate what FL would be at a given muscle length with respect to L_O_ (e.g., in trained rats for a 3-mm strain: L_O_ + 1.5 mm = 25.72 mm + 1.5 mm = 27.22 mm; FL = 27.22 mm × 0.68 = 18.51 mm; L_O_ – 1.5 mm = 25.72 mm – 1.5 mm = 24.22 mm; FL = 24.22 mm × 0.68 = 16.47 mm). Subsequently, we can estimate the SL at that muscle length by dividing FL by the SSN (e.g., at L_O_ + 1.5 mm, 18.51 mm / 6024 = 3.07 μm; at L_O_ – 1.5 mm, 16.47 mm / 6024 = 2.73 μm). Assuming the width of the sarcomere force-length relationship changed proportionately to SL, we can calculate the percentage that SL deviated from optimal SL during the whole-muscle excursion (e.g. [{3.07 – 2.93} / 2.93] × 100% = ±4.78%). We can also estimate the sarcomere shortening velocity in an excursion by multiplying the sarcomere displacement by 2 times the cycle frequency (e.g., [3.07 μm – 2.73 μm] × (1.5 Hz × 2) = 1.02 μm/s). Because optimal SL is different between groups, these velocities should also be expressed as SL/s for comparisons (e.g., 1.02 μm/s / 2.93 μm = 0.11 SL/s).

**Table 1:** Estimated sarcomere operating ranges and shortening velocities in the work loops that improved with training

| | Control (measured resting length at $L_0$: 2.81 $\mu\text{m}$) | | | Trained (measured resting length at $L_0$: 2.93 $\mu\text{m}$) | | |
| --- | --- | --- | --- | --- | --- | --- |
| Strain | SL range | Excursion from optimal SL | Shortening velocity at 1.5 Hz | SL range | Excursion from optimal SL | Shortening velocity at 1.5 Hz |
| 1 mm | 2.73-2.85 $\mu\text{m}$ | $\pm 1.42\%$ | 0.13 SL/s | 2.85-2.96 $\mu\text{m}$ | $\pm 1.02\%$ | 0.11 SL/s |
| 3 mm | 2.62-2.96 $\mu\text{m}$ | $\pm 5.34\%$ | 0.36 SL/s | 2.73-3.07 $\mu\text{m}$ | $\pm 4.78\%$ | 0.35 SL/s |
| 5 mm | 2.51-3.07 $\mu\text{m}$ | $\pm 9.25\%$ | 0.66 SL/s | 2.62-3.19 $\mu\text{m}$ | $\pm 8.87\%$ | 0.58 SL/s |

$L_0$ : optimal muscle length; SL: sarcomere length

From the estimations in Table 1, it appears that the sarcomeres of trained rats indeed shortened/lengthened relatively less from optimal SL and operated at relatively slower shortening velocities than controls at a 1.5-Hz cycle frequency with 1, 3, and 5-mm strains. Therefore, it is possible that at these cycle frequency/strain combinations, the increased SSN placed sarcomeres at more advantageous positions on the force-length and force-velocity relationships, increasing force production throughout the range of motion and contributing to greater net work output. The results from the regression analyses support these estimations. There were significant relationships between net work output and both FL and SL in only these work loops, suggesting the training-induced increases in FL and SL indeed contributed to the improved work loop performance. It appears that the increased specific maximum isometric force (i.e., improvements in force-generating capacity independent of PCSA) also contributed to the improved work loop performance, at least at the 1 and 3-mm strains, but FL and SL adaptations were more important for improved net work output at the 3 and 5-mm strains (Figure 9).

To our knowledge, one other study on animals has assessed the impact of sarcomerogenesis on work loop performance (Cox et al., 2000). After incrementally surgically stretching the rabbit latissimus dorsi for 3 weeks, they observed a 25% increase in SSN compared to controls, however, maximum work loop power output decreased by 40%, and the optimal cycle frequency shifted from 5-7 Hz to 2-3 Hz, indicating favouring for slower velocity contractions. They determined this detriment to work loop performance was related to energy loss (i.e., more negative work in passive work loops) caused by collagen accumulation, because after an additional 3 weeks of maintained stretch, collagen content decreased back toward control values, SSN increased another 5%, and work loop power output returned to normal. As the present study observed no changes in collagen content or crosslinking per unit volume of muscle, we were able to demonstrate improvements in work loop performance alongside sarcomerogenesis in the absence of intramuscular collagen accumulation.

Some studies on humans have also connected increased FL to improved mechanical work or power output. In the ankle plantar flexors, Beck et al. (2021, preprint) observed lower metabolic costs of cyclic force production in longer compared to shorter fascicle operating lengths under the same mechanical work and shortening velocity. Hinks et al. (2020) and Davidson et al. (2022) observed a 4% increase in tibialis anterior FL following 8 weeks of isometric training at a long muscle-tendon unit length, and an 11% increase in work of shortening, and 25% and 33% increases in isotonic power at loads of 10% and 50% maximum, respectively. However, they also observed increased isometric strength, pennation angle, and muscle thickness, so we can only speculate that those improvements in work and power were due to increased FL rather than strength and hypertrophy. Lastly, a computational model by Drazan et al. (2019) showed longer medial gastrocnemius fascicles produce greater force throughout the range of motion, increasing work of shortening. Further work in animals is required to elucidate whether increased FL and SSN, with constant SL, can improve net work output in cyclic contractions.

## Conclusion

The purpose of this study was to assess muscle architecture and work loop performance of the rat soleus following 4 weeks of progressively loaded weighted downhill running training. Aligning with our hypotheses, longitudinal muscle growth occurred, with a 13% increase in FL which appeared to stem from an 8% increase in SSN and a 4-5% increase in optimal SL. This longitudinal muscle growth corresponded to rightward shifts in the active and passive force-length relationships. Work loop performance improved most in the slowest cycle frequency at the 3 shortest strains. Based on regression analyses and mathematical estimations, it is feasible that these improvements in work loop performance were due to the observed longitudinal muscle growth, alongside (and to a lesser extent) improvements in force generating capacity independent of PCSA. Future research should assess the time course of SL and SSN adaptations with training, and the accompanying influence on dynamic contractile function.

## Acknowledgements

This project was supported by the Natural Sciences and Engineering Research Council of Canada (NSERC) and the Canadian Institutes of Health Research (CIHR). Infrastructure was provided by the University of Guelph start-up funding. No conflicts of interest, financial or otherwise, are declared by the authors.

## Disclosure statement

No conflicts of interest, financial or otherwise, are declared by the authors.

## Ethics statement

All procedures were approved by the Animal Care Committee of the University of Guelph.

## Data accessibility

Individual values of all supporting data are available upon request.

## Grants

This project was supported by the Natural Sciences and Engineering Research Council of Canada (NSERC) and the Canadian Institutes of Health Research (CIHR). Infrastructure was provided by the University of Guelph start-up funding.

## Author contributions

A.H., K.D.M, D.C.W, S.H.M.B., and G.A.P. conceived and designed research; A.H., K.J., and P.M. carried out animal husbandry and training; A.H. performed experiments; A.H. and K.J. analyzed data; A.H., M.V.F., S.H.M.B., and G.A.P. interpreted results of experiments; A.H. prepared figures; A.H. and G.A.P. drafted manuscript; A.H., K.J., P.M., K.D.M., M.V.F., D.C.W., S.H.M.B., and G.A.P. edited and revised manuscript; A.H., K.J., P.M., K.D.M., M.V.F., D.C.W., S.H.M.B., and G.A.P. approved final version of manuscript.

## Body weight

Figure S1 shows body weight from the start of the training period to sacrifice. In control and trained rats, body weight increased up to 17 weeks of age (week 4 of training) (all comparisons *P* < 0.01), then plateaued from 17 to 18 weeks of age (control: *P* = 0.74, trained: *P* = 0.09). While visually it appears that trained rats tended to weigh increasingly less than controls across the training period, there were no differences in body weight between trained and control rats at any weeks (*P =* 0.20-0.50). This observation strengthens comparability between the training and control groups, as it discounts differences in body weight as a confounding variable.

**Figure S1:**
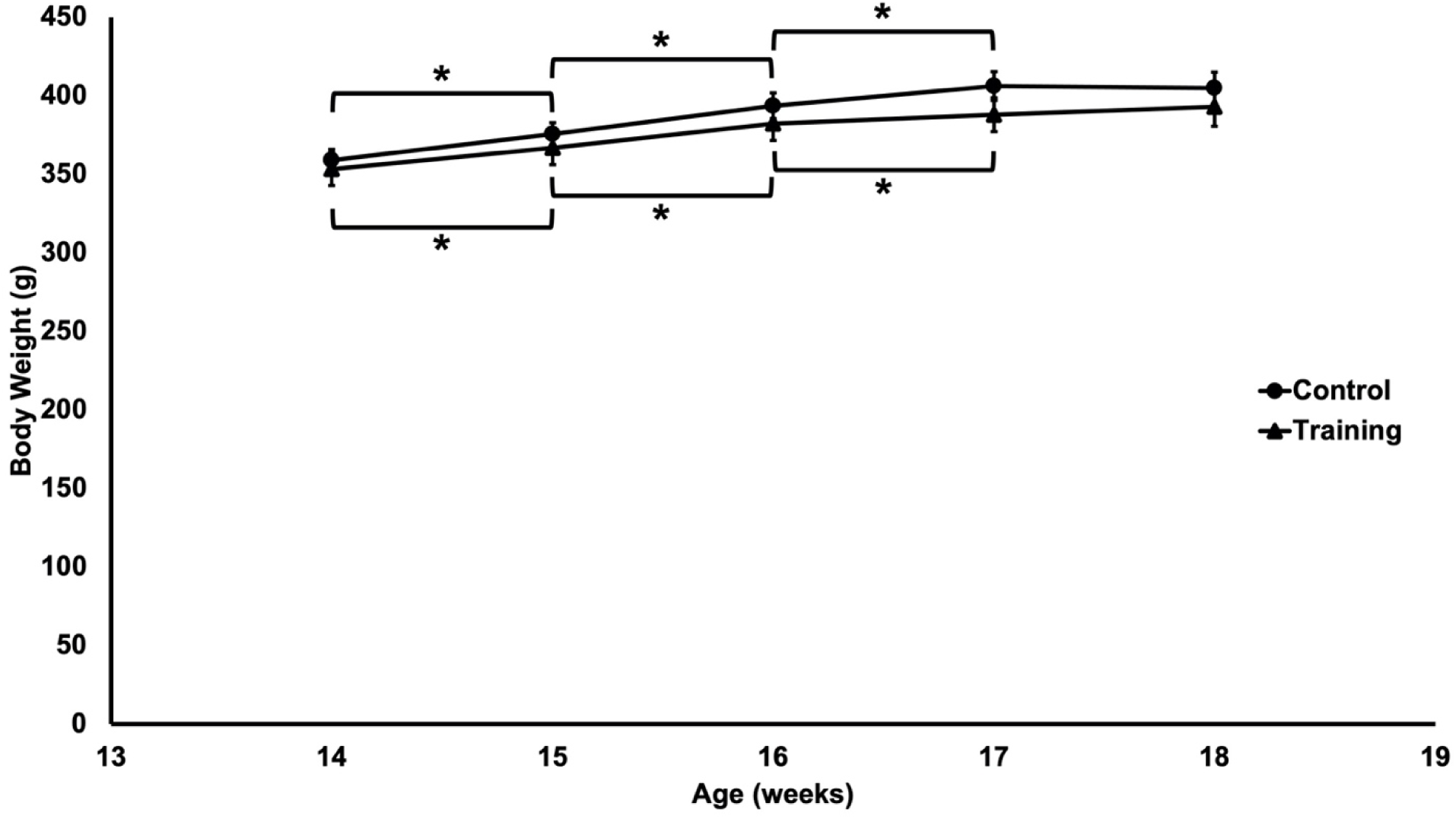
Changes in rat body weight from the start of the training period (age ∼14 weeks) to the day of sacrifice (age ∼18 weeks) (n = 18 control, n = 14 training). Data are reported as mean ± standard error. *Significant difference between time points.

**Figure S2:**
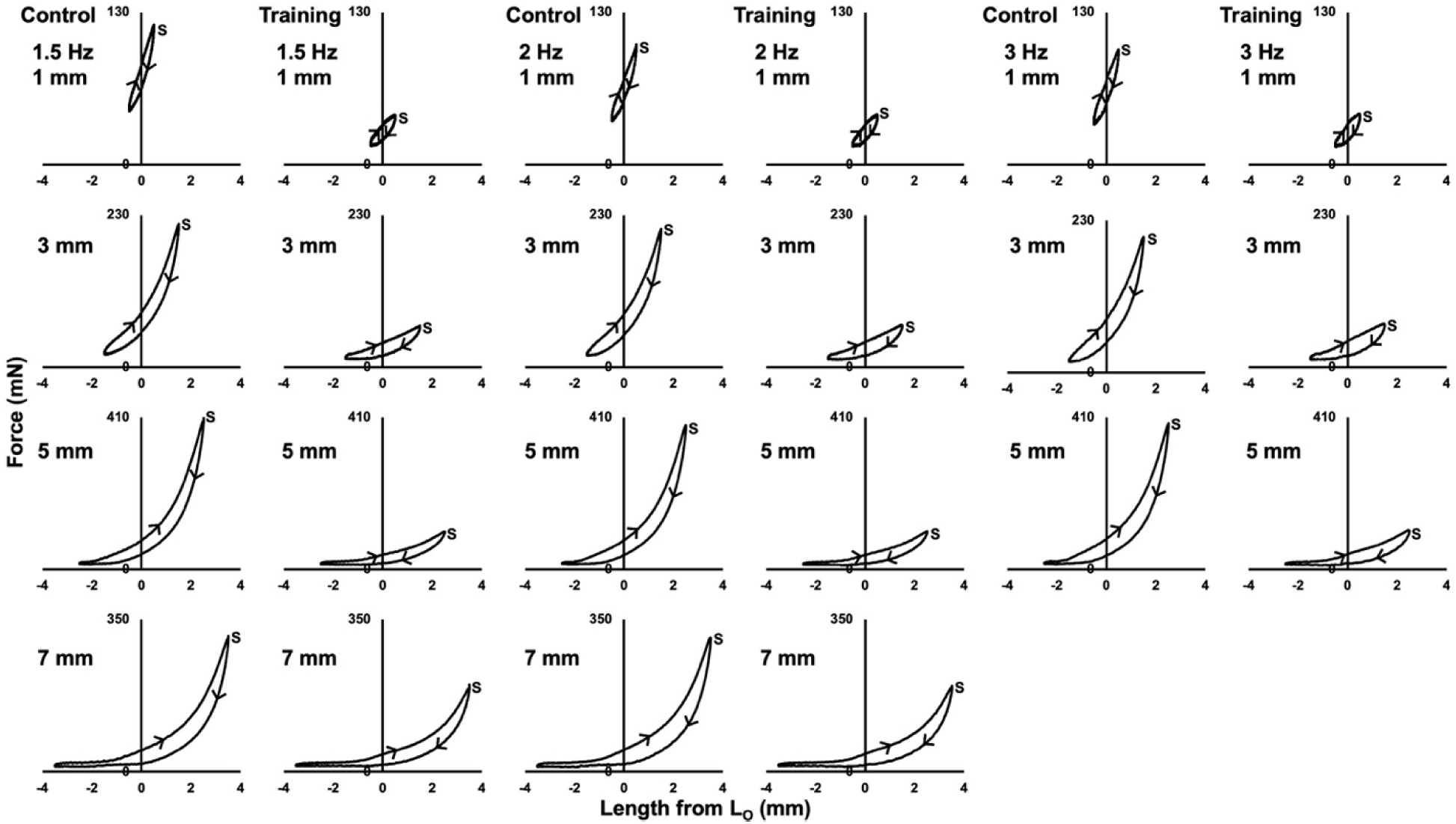
Representative passive (i.e., no stimulation) work loop traces for 1 control and 1 traine d rat across cycle frequencies of 1.5, 2, and 3 Hz, and strains of 1, 3, 5, and 7 mm. *S* indicates the start of the cycle. Arrows indicate the direction of the cycle.

**Figure S3:**
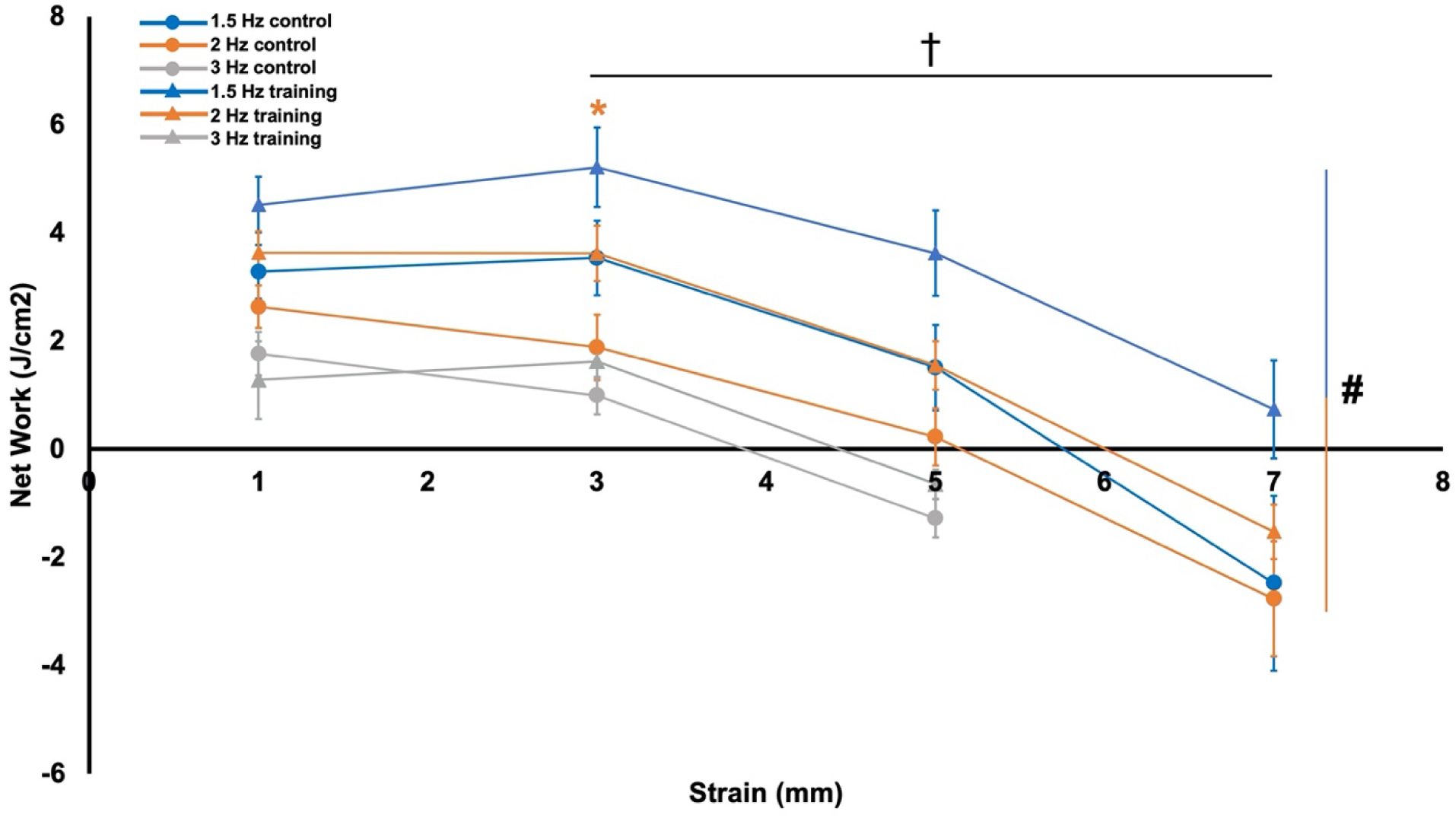
Estimated optimal net work output (i.e., work of shortening (positive) from the active work loops plus work of lengthening (negative) from the passive work loops) in control and traine d rats. Data are reported as mean ± standard error (n = 16 control, n = 12 training). *Signific ant difference (*P* < 0.05) between control and training at the colour-coded cycle frequency. #Significant difference across cycle frequencies. †Significant difference across strains.

Across both groups, the estimated optimal net work output was greater than the measured (i.e., suboptimal) net work output in the 1.5-Hz loops at 1, 3, and 5-mm strains, 2-Hz loops at 1, 3, 5, and 7-mm strains, and 3-Hz loops at 1, 3, and 5-mm strains (all *P* < 0.01), but not the 1.5-Hz loop at a 7-mm strain (*P* = 0.19). Unlike in the measured work loops, estimated optimal net work output occurred on average in the 1.5-Hz loop at a 3-mm strain in both trained and control rats. Like the suboptimal net work output, there was a group effect on estimated optimal net work output (F(1,304) = 18.51, *P* < 0.01, η_p_ ^2^ = 0.06) with trained rats producing on average 1.30 J/cm^2^ (95% CI [0.73, 1.88] more net work than controls across all work loops. There were no interactions of group × cycle frequency × strain (F(5,304) = 0.28, *P* = 0.93), group × cycle frequency (F(2,304) = 2.27, *P* = 0.11), or group × strain (F(3,304) = 0.68, *P* = 0.56), suggesting differences between groups did not depend on cycle frequency or strain. Post-hoc tests indicated specifically estimated optimal net work output was 92% greater in trained than control rats in in the 2-Hz, 3-mm work loop (F(1,26) = 4.35, *P* < 0.05, *d* = 0.83) (Figure S3).

For estimated optimal net work output, there was an effect of cycle frequency (F(2,304) = 30.56, *P* < 0.01, η_p_ ^2^ = 0.18), and strain (F(2,304) = 52.13, *P* < 0.01, η_p_ ^2^ = 0.36). Specifica lly, estimated optimal net work output also decreased with increasing cycle frequency expect for between 2 and 3 Hz (*P* = 0.40; all other comparisons *P* < 0.01), and increased with strain except for between 1 and 3 mm (*P* = 1.00; all other comparisons *P* < 0.01) (Figure S3).

## Notes

### Competing Interest Statement

The authors have declared no competing interest.

## References

Akagi, R., Hinks, A. and Power, G. A. (2020). Differential changes in muscle architecture and neuromuscular fatigability induced by isometric resistance training at short and long muscle-tendon unit lengths. J. Appl. Physiol. 129, 173–184.

Alcazar, J., Csapo, R., Ara, I. and Alegre, L. M. (2019). On the Shape of the Force-Velocity Relationship in Skeletal Muscles: The Linear, the Hyperbolic, and the Double-Hyperbolic. Front. Physiol. 10, 769.

Alder, A. B., Crawford, G. N. C. and Edwards, R. G. (1958). The growth of the muscle tibialis anterior in the normal rabbit in relation to the tension-length ratio. Proc. R. Soc. Lond. Ser. B - Biol. Sci. 148, 207–216.

Aoki, M. S., Soares, A. G., Miyabara, E. H., Baptista, I. L. and Moriscot, A. S. (2009). Expression of genes related to myostatin signaling during rat skeletal muscle longitudinal growth: Myostatin and Longitudinal Growth. Muscle Nerve 40, 992–999.

Armstrong, R. B. and Phelps, R. O. (1984). Muscle fiber type composition of the rat hindlimb. Am. J. Anat. 171, 259–272.

Baker, J. H. and Hall-Craggs, E. C. B. (1978). Changes in length of sarcomeres following tenotomy of the rat soleus muscle. Anat. Rec. 192, 55–58.

Baxter, J. R., Hullfish, T. J. and Chao, W. (2018). Functional deficits may be explained by plantarflexor remodeling following Achilles tendon rupture repair: Preliminary findings. J. Biomech. 79, 238–242.

Beck, O. N., Schroeder, J. N., Trejo., L. H., Franz, J. R. and Sawicki, G. S. (2021). Relatively Shorter Muscle Lengths Increase the Metabolic Rate of Cyclic Force Production. bioRXiv.

Bogomolovas, J., Fleming, J. R., Franke, B., Manso, B., Simon, B., Gasch, A., Markovic, M., Brunner, T., Knöll, R., Chen, J., et al. (2021). Titin kinase ubiquitination aligns autophagy receptors with mechanical signals in the sarcomere. EMBO Rep. 22,.

Brashear, S. E., Wohlgemuth, R. P., Gonzalez, G. and Smith, L. R. (2020). Passive stiffness of fibrotic skeletal muscle in mdx mice relates to collagen architecture. J. Physiol. Online ahead of print.

Butterfield, T. A. and Herzog, W. (2006). The magnitude of muscle strain does not influence serial sarcomere number adaptations following eccentric exercise. Pflüg. Arch. - Eur. J. Physiol. 451, 688–700.

Butterfield, T. A., Leonard, T. R. and Herzog, W. (2005a). Differential serial sarcomere number adaptations in knee extensor muscles of rats is contraction type dependent. J Appl Physiol 99, 7.

Butterfield, T. A., Leonard, T. R. and Herzog, W. (2005b). Differential serial sarcomere number adaptations in knee extensor muscles of rats is contraction type dependent. J. Appl. Physiol. 99, 1352–1358.

Caiozzo, V. J. and Baldwin, K. M. (1997). Determinants of work produced by skeletal muscle: potential limitations of activation and relaxation. Am. J. Physiol.-Cell Physiol. 273, C1049–C1056.

Chen, J., Mashouri, P., Fontyn, S., Valvano, M., Elliott-Mohamed, S., Noonan, A. M., Brown, S. H. M. and Power, G. A. (2020). The influence of training-induced sarcomerogenesis on the history dependence of force. J. Exp. Biol. 223, jeb218776.

Coutinho, E. L., Gomes, A. R. S., França, C. N., Oishi, J. and Salvini, T. F. (2004). Effect of passive stretching on the immobilized soleus muscle fiber morphology. Braz. J. Med. Biol. Res. 37, 1853–1861.

Cox, V. M., Williams, P. E., Wright, H., James, R. S., Gillott, K. L., Young, I. S. and Goldspink, D. F. (2000). Growth induced by incremental static stretch in adult rabbit latissimus dorsi muscle. Exp Physiol 85, 193–202.

Davidson, B., Hinks, A., Dalton, B. H., Akagi, R. and Power, G. A. (2022). Power attenuation from restricting range of motion is minimized in subjects with fast RTD and following isometric training. J. Appl. Physiol. 132, 497–510.

Davis, J. F., Khir, A. W., Barber, L., Reeves, N. D., Khan, T., DeLuca, M. and Mohagheghi, A. A. (2020). The mechanisms of adaptation for muscle fascicle length changes with exercise: Implications for spastic muscle. Med. Hypotheses 144, 110199.

De Jaeger, D., Joumaa, V. and Herzog, W. (2015). Intermittent stretch training of rabbit plantarflexor muscles increases soleus mass and serial sarcomere number. J. Appl. Physiol. 118, 1467–1473.

De Koning, J. J., Van Der Molen, H. F., Woittiez, R. D. and Huijing, P. A. (1987). Functional characteristics of rat gastrocnemius and tibialis anterior muscles during growth. J. Morphol. 194, 75–84.

Drazan, J. F., Hullfish, T. J. and Baxter, J. R. (2019). Muscle structure governs joint function: linking natural variation in medial gastrocnemius structure with isokinetic plantar flexor function. Biol. Open bio.048520.

Farup, J., Kjølhede, T., Sørensen, H., Dalgas, U., Møller, A. B., Vestergaard, P. F., Ringgaard, S., Bojsen-Møller, J. and Vissing, K. (2012). Muscle Morphological and Strength Adaptations to Endurance Vs. Resistance Training. J. Strength Cond. Res. 26, 398–407.

Fitts, R. H., McDonald, K. S. and Schluter, J. M. (1991). The determinants of skeletal muscle force and power: Their adaptability with changes in activity pattern. J. Biomech. 24, 111– 122.

Franchi, M. V., Ruoss, S., Valdivieso, P., Mitchell, K. W., Smith, K., Atherton, P. J., Narici, M. V. and Flück, M. (2018). Regional regulation of focal adhesion kinase after concentric and eccentric loading is related to remodelling of human skeletal muscle. Acta Physiol. 223, e13056.

Gans, C. and Bock, W. J. (1965). The functional significance of muscle architecture--a theoretical analysis. Ergeb Anat Entwicklungsgesch 38, 115–42.

Gans, C. and de Vree, F. (1987). Functional bases of fiber length and angulation in muscle. J. Morphol. 192, 63–85.

Gillies, A. R. and Lieber, R. L. (2011). Structure and function of the skeletal muscle extracellular matrix: Skeletal Muscle ECM. Muscle Nerve 44, 318–331.

Gokhin, D. S., Dubuc, E. A., Lian, K. Q., Peters, L. L. and Fowler, V. M. (2014). Alterations in thin filament length during postnatal skeletal muscle development and aging in mice. Front. Physiol. 5,.

Gordon, A. M., Huxley, A. F. and Julian, F. J. (1966a). The variation in isometric tension with sarcomere length in vertebrate muscle fibres. J. Physiol. 184, 170–192.

Gordon, A. M., Huxley, A. F. and Julian, F. J. (1966b). Tension development in highly stretched vertebrate muscle fibres. J. Physiol. 184, 143–169.

Griffiths, R. I. (1991). Shortening of muscle fibres during stretch of the active cat medial gastrocnemius muscle: the role of tendon compliance. J. Physiol. 436, 219–236.

Han, X.-Y., Wang, W., Koskinen, S. O. A., Kovanen, V., Takala, T. E. S., Komulainen, J., Vihko, V. and Trackman, P. C. (1999). Increased mRNAs for procollagens and key regulating enzymes in rat skeletal muscle following downhill running. Pflüg. Arch. Eur. J. Physiol. 437, 857–864.

Haun, C. T., Vann, C. G., Roberts, B. M., Vigotsky, A. D., Schoenfeld, B. J. and Roberts, M. D. (2019). A Critical Evaluation of the Biological Construct Skeletal Muscle Hypertrophy: Size Matters but So Does the Measurement. Front. Physiol. 10, 247.

Heinemeier, K. M., Olesen, J. L., Haddad, F., Langberg, H., Kjaer, M., Baldwin, K. M. and Schjerling, P. (2007). Expression of collagen and related growth factors in rat tendon and skeletal muscle in response to specific contraction types: Collagen and TGF-β-1 expression in exercised tendon and muscle. J. Physiol. 582, 1303–1316.

Herbert, R. D. and Balnave, R. J. (1993). The effect of position of immobilisation on resting length, resting stiffness, and weight of the soleus muscle of the rabbit. J. Orthop. Res. 11, 358–366.

Herbert, R. D. and Gandevia, S. C. (2019). The passive mechanical properties of muscle. J. Appl. Physiol. 126, 1442–1444.

Herring, S. W., Grimm, A. F. and Grimm, B. R. (1984). Regulation of sarcomere number in skeletal muscle: A comparison of hypotheses. Muscle Nerve 7, 161–173.

Herzog, W. and Fontana, H. de B. (2021). Does eccentric exercise stimulate sarcomerogenesis? J. Sport Health Sci. S2095254621001083.

Heslinga, J. W. and Huijing, P. A. (1993). Muscle length-force characteristics in relation to muscle architecture: a bilateral study of gastrocnemius medialis muscles of unilaterally immobilized rats. Eur. J. Appl. Physiol. 66, 289–298.

Hinks, A., Davidson, B., Akagi, R. and Power, G. A. (2021). Influence of isometric training at short and long muscle-tendon unit lengths on the history dependence of force. Scand. J. Med. Sci. Sports 31, 325–338.

Honda, Y., Tanaka, M., Tanaka, N., Sasabe, R., Goto, K., Kataoka, H., Sakamoto, J., Nakano, J. and Okita, M. (2018). Relationship between extensibility and collagen expression in immobilized rat skeletal muscle. Muscle Nerve 57, 672–678.

Huijing, P. A. (1999). Muscle as a collagen fiber reinforced composite: a review of force transmission in muscle and whole limb. J. Biomech. 32, 329–345.

Hyldahl, R. D. and Hubal, M. J. (2014). Lengthening our perspective: Morphological, cellular, and molecular responses to eccentric exercise: Exercise-Induced Muscle Damage. Muscle Nerve 49, 155–170.

Jahromi, S. S. and Charlton, M. P. (1979). Transverse sarcomere splitting. A possible means of longitudinal growth in crab muscles. J. Cell Biol. 80, 736–742.

Jakubiec-Puka, A. and Carraro, U. (1991). Remodelling of the contractile apparatus of striated muscle stimulated electrically in a shortened position. J. Anat. 178, 83–100.

James, R. S., Altringham, J. D. and Goldspink, D. F. (1995). THE MECHANICAL PROPERTIES OF FAST AND SLOW SKELETAL MUSCLES OF THE MOUSE IN RELATION TO THEIR LOCOMOTORY FUNCTION. J. Exp. Biol. 198, 491–502.

Jorgenson, K. W., Phillips, S. M. and Hornberger, T. A. (2020). Identifying the Structural Adaptations that Drive the Mechanical Load-Induced Growth of Skeletal Muscle: A Scoping Review. Cells 9, 1658.

Josephson, R. K. (1999). Dissecting muscle power output. J. Exp. Biol. 202, 3369–3375.

Josephson, R. K. and Stokes, D. R. (1989). STRAIN,MUSCLE LENGTH ANDWORK OUTPUT IN A CRAB MUSCLE. J. Exp. Biol. 145, 45–61.

Karpakka, J. A., Pesola, M. K. and Takala Timo E. S. (1992). Thee effects of anabolic steroids on collagen synthesis in rat skeletal muscle and tendon. Am. J. Sports Med. 20, 262–266.

Kjær, M. (2004). Role of Extracellular Matrix in Adaptation of Tendon and Skeletal Muscle to Mechanical Loading. Physiol. Rev. 84, 649–698.

Koh, T. J. (1995). Do adaptations in serial sarcomere number occur with strength training? Hum. Mov. Sci. 14, 61–77.

Koh, T. J. and Tidball, J. G. (1999). Nitric oxide synthase inhibitors reduce sarcomere addition in rat skeletal muscle. J. Physiol. 519, 189–196.

Lichtwark, G. A. and Barclay, C. J. (2010). The influence of tendon compliance on muscle power output and efficiency during cyclic contractions. J. Exp. Biol. 213, 707–714.

Lieber, R. L. and Ward, S. R. (2011). Skeletal muscle design to meet functional demands. Philos. Trans. R. Soc. B Biol. Sci. 366, 1466–1476.

Lieber, R. L., Yeh, Y. and Baskin, R. J. (1984). Sarcomere length determination using laser diffraction. Effect of beam and fiber diameter. Biophys. J. 45, 1007–1016.

Lindstedt, S. L., McGlothlin, T., Percy, E. and Pifer, J. (1998). Task-specific design of skeletal muscle: balancing muscle structural composition. Comp. Biochem. Physiol. B Biochem. Mol. Biol. 120, 35–40.

Łochyński, D., Kaczmarek, D., Grześkowiak, M., Majerczak, J., Podgórski, T. and Celichowski, J. (2021). Motor Unit Force Potentiation and Calcium Handling Protein Concentration in Rat Fast Muscle After Resistance Training. Front. Physiol. 12, 652299.

Lutz, G. J. and Rome, L. C. (1994). Built for Jumping: The Design of the Frog Muscular System. Sci. New Ser. 263, 370–372.

Lynn, R., Talbot, J. A. and Morgan, D. L. (1998). Differences in rat skeletal muscles after incline and decline running. J. Appl. Physiol. 85, 98–104.

Macadam, P., Cronin, J. B. and Feser, E. H. (2019). Acute and longitudinal effects of weighted vest training on sprint-running performance: a systematic review. Sports Biomech. 1–16.

Morais, G. P., da Rocha, A. L., Neave, L. M., de A. Lucas, G., Leonard, T. R., Carvalho, A., da Silva, A. S. R. and Herzog, W. (2020). Chronic uphill and downhill exercise protocols do not lead to sarcomerogenesis in mouse skeletal muscle. J. Biomech. 98, 109469.

Narici, M., Franchi, M. and Maganaris, C. (2016). Muscle structural assembly and functional consequences. J. Exp. Biol. 219, 276–284.

Nicolopoulos-Stournaras, S. and Iles, J. F. (1983). Hindlimb muscle activity during locomotion in the rat (Rattus norvegicus) (Rodentia: Muridae). J. Zool. 203, 427–440.

Noonan, A. M., Mashouri, P., Chen, J., Power, G. A. and Brown, S. H. M. (2020). Training Induced Changes to Skeletal Muscle Passive Properties Are Evident in Both Single Fibers and Fiber Bundles in the Rat Hindlimb. Front. Physiol. 11, 907.

Ochi, E., Nakazato, K. and Ishii, N. (2007). Effects of Eccentric Exercise on Joint Stiffness and Muscle Connectin (Titin) Isoform in the Rat Hindlimb. J. Physiol. Sci. 57, 1–6.

Ranatunga, K. (2018). Temperature Effects on Force and Actin–Myosin Interaction in Muscle: A Look Back on Some Experimental Findings. Int. J. Mol. Sci. 19, 1538.

Roberts, T. J. (2016). Contribution of elastic tissues to the mechanics and energetics of muscle function during movement. J. Exp. Biol. 219, 266–275.

Rome, L. C. (1994). The mechanical design of the fish muscular system. Mech. Physiol. Anim. Swim. 75–98.

Roots, H. and Ranatunga, K. W. (2008). An analysis of the temperature dependence of force, during steady shortening at different velocities, in (mammalian) fast muscle fibres. J. Muscle Res. Cell Motil. 29, 9–24.

Roy, R. R., Meadows, I. D., Baldwin, K. M. and Edgerton, V. R. (1982). Functional significance of compensatory overloaded rat fast muscle. J. Appl. Physiol. 52, 473–478.

Roy, R. R., Hutchison, D. L., Pierotti, D. J., Hodgson, J. A. and Edgerton, V. R. (1991). EMG patterns of rat ankle extensors and flexors during treadmill locomotion and swimming. J. Appl. Physiol. 70, 2522–2529.

Roy, R. R., Zhong, H., Monti, R. J., Vallance, K. A. and Edgerton, V. R. (2002). Mechanical properties of the electrically silent adult rat soleus muscle. Muscle Nerve 26, 404–412.

Salzano, M. Q., Cox, S. M., Piazza, S. J. and Rubenson, J. (2018). American Society of Biomechanics Journal of Biomechanics Award 2017: High-acceleration training during growth increases optimal muscle fascicle lengths in an avian bipedal model. J. Biomech. 80, 1–7.

Schaeffer, P. J. and Lindstedt, S. L. (2013). How Animals Move: Comparative Lessons on Animal Locomotion. Compr. Physiol. 3, 289–314.

Schilder, R. J., Kimball, S. R., Marden, J. H. and Jefferson, L. S. (2011). Body weight-dependent troponin T alternative splicing is evolutionarily conserved from insects to mammals and is partially impaired in skeletal muscle of obese rats. J. Exp. Biol. 214, 1523–1532.

Shrager, J. B., Kim, D.-K., Hashmi, Y. J., Stedman, H. H., Zhu, J., Kaiser, L. R. and Levine, S. (2002). Sarcomeres Are Added in Series to Emphysematous Rat Diaphragm After Lung Volume Reduction Surgery. Chest 121, 210–215.

Soares, A. G., Aoki, M. S., Miyabara, E. H., DeLuca, C. V., Ono, H. Y., Gomes, M. D. and Moriscot, A. S. (2007). Ubiquitin-ligase and deubiquitinating gene expression in stretched rat skeletal muscle. Muscle Nerve 36, 685–693.

Spletter, M. L., Barz, C., Yeroslaviz, A., Zhang, X., Lemke, S. B., Bonnard, A., Brunner, E., Cardone, G., Basler, K., Habermann, B. H., et al. (2018). A transcriptomics resource reveals a transcriptional transition during ordered sarcomere morphogenesis in flight muscle. eLife 7, e34058.

Sugama, S., Tachino, K. and Haida, N. (1999). Effect of Immobilization on Solubility of Soleus and Gastrocnemius Muscle Collagen. Biochemical Studies on Collagen from Soleus and Gastrocnemius Muscles of Rat. J. Jpn. Phys. Ther. Assoc. 2, 25–29.

Swoap, S. J., Caiozzo, V. J. and Baldwin, K. M. (1997). Optimal shortening velocities for in situ power production of rat soleus and plantaris muscles. Am. J. Physiol.-Cell Physiol. 273, C1057–C1063.

Tabary, J. C., Tabary, C., Tardieu, C., Tardieu, G. and Goldspink, G. (1972). Physiological and structural changes in the cat’s soleus muscle due to immobilization at different lengths by plaster casts*. J. Physiol. 224, 231–244.

Takahashi, M., Ward, S. R., Fridén, J. and Lieber, R. L. (2012). Muscle excursion does not correlate with increased serial sarcomere number after muscle adaptation to stretched tendon transfer: FUNCTIONAL ADAPTATION AFTER TENDON TRANSFER. J. Orthop. Res. 30, 1774–1780.

Turrina, A., Martínez-González, M. A. and Stecco, C. (2013). The muscular force transmission system: Role of the intramuscular connective tissue. J. Bodyw. Mov. Ther. 17, 95–102.

Walker, S. M. and Schrodt, G. R. (1974). I segment lengths and thin filament periods in skeletal muscle fibers of the rhesus monkey and the human. Anat. Rec. 178, 63–81.

Ward, S. R. and Lieber, R. L. (2005). Density and hydration of fresh and fixed human skeletal muscle. J. Biomech. 38, 2317–2320.

Wickiewicz, T. L., Roy, R. R., Powell, P. L., Perrine, J. J. and Edgerton, V. R. (1984). Muscle architecture and force-velocity relationships in humans. J. Appl. Physiol. 57, 435–443.

Widrick, J. J., Maddalozzo, G. F., Hu, H., Herron, J. C., Iwaniec, U. T. and Turner, R. T. (2008). Detrimental effects of reloading recovery on force, shortening velocity, and power of soleus muscles from hindlimb-unloaded rats. Am. J. Physiol.-Regul. Integr. Comp. Physiol. 295, R1585–R1592.

Williams, P. E. and Goldspink, G. (1973). The effect of immobilization on the longitudinal growth of striated muscle fibres. J. Anat. 116, 45–55.

Williams, P. E. and Goldspink, G. (1978). Changes in sarcomere length and physiological properties in immobilized muscle. J. Anat. 127, 459–468.

Woittiez, R. D., Baan, G. C., Huijing, P. A. and Rozendal, R. H. (1985). Functional characteristics of the calf muscles of the rat. J. Morphol. 184, 375–387.

Woittiez, R. D., Heerkens, Y. F., Huijing, P. A., Rijnsburger, W. H. and Rozendal, R. H. (1986). Functional morphology of the M. Gastrocnemius medialis of the rat during growth. J. Morphol. 187, 247–258.

Wong, T. S. and Booth, F. W. (1988). Skeletal muscle enlargement with weight-lifting exercise by rats. J. Appl. Physiol. 65, 950–954.

Xu, H., Ren, X., Lamb, G. D. and Murphy, R. M. (2018). Physiological and biochemical characteristics of skeletal muscles in sedentary and active rats. J. Muscle Res. Cell Motil. 39, 1–16.

Zhu, W. G., Hibbert, J. E., Lin, K.-H., Steinert, N. D., Lemens, J. L., Jorgenson, K. W., Newman, S. M., Lamming, D. W. and Hornberger, T. A. (2021). Weight Pulling: A Novel Mouse Model of Human Progressive Resistance Exercise. Cells 10, 2549.

Zimmerman, S. D., McCormick, R. J., Vadlamudi, R. K. and Thomas, D. P. (1993). Age and training alter collagen characteristics in fast- and slow-twitch rat limb muscle. J. Appl. Physiol. 75, 1670–1674.

Zöllner, A. M., Abilez, O. J., Böl, M. and Kuhl, E. (2012). Stretching Skeletal Muscle: Chronic Muscle Lengthening through Sarcomerogenesis. PLoS ONE 7, e45661.

